# Episodic Transport of Protein Aggregates Achieves a Positive Size Selectivity in Aggresome Formation

**DOI:** 10.1101/2024.08.06.606767

**Authors:** Rui Fang, Luolan Bai, Boyan Li, Kevin Dong, Joao A. Paulo, Mengying Zhou, Yi-Chi Chu, Yuyu Song, Michael Y. Sherman, Steven Gygi, Christine M. Field, Timothy J. Mitchison, Ying Lu

**Affiliations:** Department of Systems Biology, Harvard Medical School, Boston, MA, USA; Department of Cell Biology, Harvard Medical School, Boston, MA, USA; Department of Neurology, Massachusetts General Hospital, Boston, MA, USA; Department of Molecular Biology, Ariel University, Ariel, Israel; School of Life Sciences, Westlake University, Hang Zhou, China

## Abstract

Eukaryotic cells direct toxic misfolded proteins to various protein quality control pathways based on their chemical features and aggregation status. Aggregated proteins are targeted to selective autophagy or specifically sequestered into the “aggresome,” a perinuclear inclusion at the microtubule-organizing center (MTOC). However, the mechanism for selectively sequestering protein aggregates into the aggresome remains unclear. To investigate aggresome formation, we reconstituted MTOC-directed aggregate transport in *Xenopus laevis* egg extract using AgDD, a chemically inducible aggregation system. High-resolution single-particle tracking revealed that dynein-mediated transport of aggregates was highly episodic, with average velocity positively correlated with aggregate size. Our mechanistic model suggests that the recurrent formation of the dynein transport complex biases larger aggregates towards the active transport state, compensating for the slowdown due to viscosity. Both episodic transport and positive size selectivity are specifically associated with aggresome-dynein adaptors. Coupling conventional dynein-activating adaptors to the aggregates perturbs aggresome formation and reverses size selectivity.

## Introduction

Accumulation of damaged or misfolded proteins leads to cytotoxicity and underlies a variety of human disorders, such as neurodegenerative diseases and type-II diabetes^1^. To maintain protein homeostasis, eukaryotic cells possess multiple protein quality control (PQC) systems that operate complementarily to eliminate abnormal proteins with diverse chemical and physical properties.

The proteasome system is responsible for degrading most soluble substrates. Proteins that aggregate into higher-order structures, such as oligomers and aggregates, are often resistant to proteasomal degradation and can even inhibit proteasome activities, necessitating their clearance through alternative PQC pathways, such as selective autophagy or aggresome sequestration. The aggresome is a membrane-less organelle localized at the microtubule-organizing center (MTOC)^2,3^. Aggresome formation represents a general cellular defense mechanism that is activated when the production of aggregation-prone proteins exceeds the degradation capacity^4^.

Apart from its protective role, the aggresome may serve as a precursor to some pathological inclusions observed in liver and neurodegenerative diseases, and a similar process has been suggested in inflammasome formation and viral replication^5–9^. The mechanism by which misfolded proteins are selectively targeted into the aggresome remains poorly understood.

Formation of the aggresome typically initiates with the nucleation of aggregation-prone proteins into small aggregates in the peripheral cytoplasm, which are termed “pre-aggresome particles,” followed by the transport of these aggregates along microtubules to the perinuclear region by the dynein motor complex^10–15^. A variety of enzymes, molecular chaperones, and adaptor proteins have been discovered to regulate this process. For instance, HDAC6, SQSTM1/p62, and the Hsp70/BAG3/14-3-3 complex recognize misfolded proteins and bridge them with the dynein machinery^3^. HDAC6 interacts with the ubiquitin (Ub) or Ub chain on substrates, and bridges them with p150^Glued^, a core component in the dynactin complex responsible for dynein activation, while SQSTM1 may interact with dynein intermediate chains (DIC) and recruit the substrates via the associated Lys63-linked Ub chains^14,16,17^. Members of the 14-3-3 family also interact with DIC; their recognition of misfolded protein is independent of Ub and presumably mediated by the interacting partner Hsp70^14^. For simplicity, we refer to these factors that link protein substrates with the dynein motor as aggresome adaptors. Different aggresome adaptors can work concurrently in the same cell to promote the sequestration of protein aggregates^3,14,18^, while the substrate specificity of each aggresome adaptor has not been fully understood. Besides the aforementioned aggresome adaptors, additional factors, such as ataxin-3, that interact with both dynein and misfolded proteins, have been reported to regulate aggresome formation^12,15^.

Unlike protein aggregates, the transport of dynein’s conventional cargoes, such as cellular vesicles and organelles, is mainly mediated by the activating adaptors, including BICD2, HOOK2, and HOOK3, which link these cargoes to the dynein motor. Besides substrate selection, the activating adaptors also stabilize dynein-dynactin interaction and activate cargo transport by stimulating dynein’s speed and processivity^19,20^. All activating adaptors contain a coiled-coil domain that interacts extensively with both dynein and dynactin^20^. However, aggresome adaptors lack the coiled-coil domain and may exhibit weak interactions with dynein, with HDAC6 as an example^13^. The functional implications for employing structurally distinct adaptors for aggregate transport have not been explored.

Directing misfolded proteins into appropriate PQC pathways requires selectivity based on the aggregation status (or size) of the protein. Protein aggregates appear to be preferentially sorted into the autophagy pathway or loaded onto the dynein motor for aggresome sequestration, over soluble misfolded proteins^21^. HDAC6’s BUZ domain interacts with the free C-terminus of Ub or Ub chain that is generated through substrate deubiquitylation. This free-Ub specificity may allow HDAC6 to interact with protein aggregates that contain trapped Ub^22^. However, some aggregation-prone proteins, such as GFP-250 and synphilin-1, are sequestered by Ub-independent aggresome pathways, and their aggresome formation is still preceded by substrate nucleation^10–12,14,23^. These observations suggest that ubiquitylation is not instrumental for aggregate selection, and a general feature of protein aggregates, e.g., their size, may guide this process. Contrary to the requirement to target aggregated proteins, dynein-mediated transport of conventional cargoes typically favors the movement of small objects, due to cytosolic viscous friction that increases with the cargo size^24–29^. How this property of transport is reconciled with the desired specificity for protein aggregates during aggresome formation is unclear. Adding to this specificity puzzle, aggresome adaptors Hsp70 and SQSTM1 are involved in diverse biological processes, including delivering soluble substrates for proteasomal degradation^30,31^. How the same factors can participate in PQC pathways with distinct substrate specificities remains unclear.

To elucidate the mechanism by which the aggresome pathways recognize the aggregated forms of misfolded proteins, we used a chemically inducible aggregation-prone protein, AgDD, and reconstituted the process of MTOC-directed transport of protein aggregates in both live cells and *Xenopus laevis* egg extract (XE). XE is one of the least perturbed cell-free systems available, and is used extensively to recapitulate intracellular dynamic structures and events^32^. Our reconstitution system successfully recapitulated the key features of protein aggregate transport in live cells, including the role of previously identified aggresome adaptors, and allowed us to gain insights into the mechanism of an unusual positive size selectivity (PSS) where larger aggregates are transported more efficiently during aggresome formation. This transport feature is specifically associated with aggresome adaptors and distinct from the transport of other cargoes, like cellular vesicles and organelles, where a negative size selectivity (NSS) is usually observed^24–28^. Through high-resolution single-particle tracking, we observed that protein aggregates exhibited “stop-and-go” movements with low average velocities. Interestingly, larger aggregates, despite having lower instantaneous velocities, experienced shorter pauses and hence achieved a higher average transport velocity. Our mechanistic model suggests that rapid disassembly and reformation of the aggregate-dynein-microtubule complex during episodic transport biases larger aggregates towards the active transport mode and quantitatively accounts for the observed PSS in aggresome formation.

## Results

### Protein aggregates are selectively targeted to the aggresome

To confirm that protein aggregates are preferentially targeted for aggresome sequestration, we employed AgDD-sfGFP (AgDD) as a model for aggregation-prone proteins. AgDD consists of an FKBP-based destabilization domain (DD) fused with a short hydrophobic peptide at the N-terminus^33^. The DD degron domain can be stabilized by the ligand Shield-1. Anti-GFP immunoblotting suggests that AgDD was stably expressed as a full-length protein in the cell lines (Fig. S1B) and depleting Shield-1 from the culture medium led to rapid misfolding and aggregation of AgDD into cytoplasmic aggregates (Fig. 1A, S1A; Supp. Video 1). These initial AgDD aggregates were then actively transported towards the nucleus to form a perinuclear aggresome-like punctum that colocalized with the centrosomal component pericentrin and was surrounded by a cage-like structure of vimentin, which are commonly utilized as markers of aggresome formation (Fig. 1B)^4^. The presence of AgDD aggresome did not significantly affect cell growth^33,34^, nor did it strongly perturb the cell-cycle phase durations as indicated by the localization of fluorescently labeled Histone H2A (Fig. S2), which greatly simplified data interpretation.

**Figure 1.**
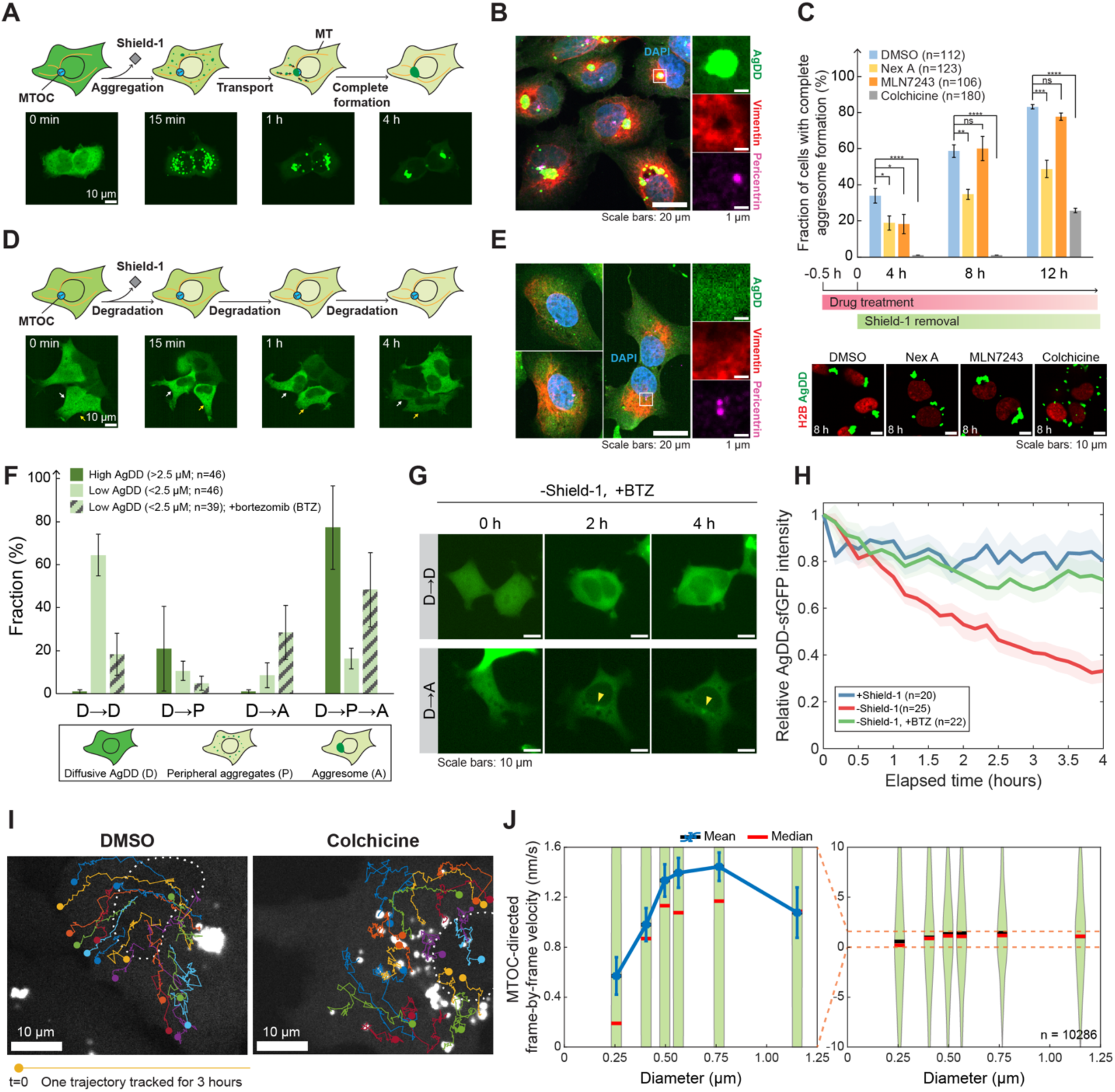
AgDD aggregates are selectively transported into the aggresome. **(A)** Schematic and representative live-cell confocal images of AgDD aggresome formation. 10 μM FKBP(F36V) was added to the culture of HEK293T cells stably expressing AgDD-sfGFP (AgDD) to deplete Shield-1. **(B)** Colocalization of AgDD with aggresome markers. AgDD (green)-expressing U2OS cells were fixed and stained with anti-vimentin (red) and anti-pericentrin (magenta) antibodies, 4 hours after Shield-1 removal. DNA was stained with DAPI (blue). **(C)** Effects of drug treatment on aggresome formation. HEK293T cells stably expressing AgDD and H2B-mCherry were treated with 20 μM Nexturastat A, 10 μM MLN7243, or 1 μg/mL colchicine for 30 minutes before Shield-1 removal to induce aggresome formation, and imaged with a spinning-disk confocal microscope. The fraction of cells that had completed aggresome formation, indicated by the sequestration of all peripheral aggregates to a single perinuclear punctum, was determined at the indicated time points. Error bars represent the standard error of the mean (SEM) over 4 randomly chosen fields of view (FOVs); n: number of cells analyzed. P-values were determined by an unpaired two-tailed Student’s *t*-test (ns: *p* > 0.05; \**p* ≤ 0.05; \*\**p* ≤ 0.01; \*\*\**p* ≤ 0.001; \*\*\*\**p* ≤ 0.0001). Below: representative images under each condition with AgDD (green) and H2B (red) channels, 8 hours after Shield-1 removal. **(D)** Schematic and live-cell confocal images as in **A** but showing cells without detectable aggresome formation after Shield-1 removal. GFP display range was adjusted for presentation. Representative images of aggresome-free cells co-stained with anti-vimentin and anti-pericentrin antibodies are shown in **E**. **(F)** Correlation between AgDD aggregation and aggresome formation. AgDD-expressing HEK293T cells were induced by Shield-1 removal with or without cotreatment of 1 μM bortezomib (BTZ) and subjected to time-lapse imaging at 10 minutes per frame for 4 hours. Randomly-chosen cells under each condition were classified according to how AgDD changed localization during the timelapse. D: diffusive AgDD signal; P: peripheral AgDD aggregates; A: AgDD aggresome. Cells with high (>2.5 μM) and low (< 2.5 μM) initial AgDD levels were counted separately. Error bars represent SEM over 4 FOVs; n: number of cells analyzed. Representative images are shown in **G**, with aggresomes indicated by arrowheads. (**H**) Degradation of diffusive AgDD upon Shield-1 removal. The diffusive AgDD signal was determined in the aggresome-free cells (“D→D” in **F**) during the timelapse, normalized to the initial AgDD intensity right after Shield-1 removal (t=0). Line shading represents SEM over n cells analyzed. **(I)** Live-cell confocal images of U2OS cells 3 hours after Shield-1 removal, overlaid with trajectories of individual AgDD aggregates whose initial positions were marked with filled circles. Cells were treated with 1 μg/mL colchicine or DMSO for 30 minutes before aggresome induction and imaged once per minute. Nuclear contours were marked with white dashed lines. **(J)** The relationship between the MTOC-directed velocity and the diameter of AgDD aggregates. The MTOC-directed velocities of individual aggregates were determined frame-by-frame, and correlated with the aggregate diameter at the same time point in a violin plot (n=10286 velocity-diameter pairs from 4 cells). Positive values of velocity represent movement towards the MTOC. Error bars in the zoom-in plot (left) represent the SEM within each size group.

Similar to other aggregation-prone proteins, AgDD requires an intact microtubule network to localize to the aggresome. Colchicine, a microtubule destabilizer, impeded the perinuclear accumulation of misfolded AgDD, resulting in multiple small aggregates accumulating in the peripheral cytoplasm (Fig. 1C). The HDAC6 inhibitor, Nexturastat A, delayed the completion of AgDD aggresome formation marked by all peripheral AgDD aggregates being sequestered into a unique perinuclear punctum, but did not completely block it, suggesting that the deacetylase activity of HDAC6 plays some role in the process as previously reported (Fig. 1C)^13^. Inhibiting the ubiquitin-activating enzyme (E1) activity with MLN7243 did not alter the timing of perinuclear sequestration of AgDD, suggesting that substrate ubiquitylation is not essential for AgDD aggresome formation (Fig. 1C, S1C).

The perinuclear deposition of misfolded AgDD correlates with, and potentially requires, its prior nucleation in the peripheral cytoplasm. To test the connection between AgDD nucleation and aggresome formation, we studied the fate of misfolded AgDD in either diffuse or aggregated states, leveraging the natural variation of AgDD expression levels in the cell population. The molar concentrations of AgDD in individual cells were determined based on calibration using purified GFP. As with many aggregation-prone proteins, the tendency of AgDD to aggregate increased with its concentration. Cells constitutively expressing a low level of AgDD did not show detectable aggregates upon Shield-1 removal (Fig. 1D, 1E). We classified the cells according to the changes in AgDD localization in time-lapse and found that cells without detectable AgDD aggregation also lacked a perinuclear accumulation of AgDD (Fig. 1F).

However, AgDD misfolding still occurred in these aggregation-free cells, as indicated by the proteasome-dependent rapid decrease of the diffusive GFP signal which could be stabilized by the proteasome inhibitor, bortezomib (BTZ) (Fig. 1H). BTZ treatment is known to induce nonspecific protein aggregation and aggresome formation^35^. Upon proteasome inhibition, most of these low-AgDD cells formed an aggresome-like perinuclear punctum within four hours after Shield-1 removal, though the intensity of the puncta appeared weak (Fig. 1F, 1G). This result excludes the possibility that these low-AgDD cells were inherently defective in aggresome formation and suggests a role for AgDD aggregation in facilitating its sequestration in the aggresome.

To quantitatively compare the efficiency of aggresome formation under different conditions, we defined the “aggresome enrichment factor” as the ratio of the maximum fluorescence intensity of pixels within the perinuclear region to the average intensity of the cell before Shield-1 removal. Although this factor for BTZ-induced aggresome in low-AgDD cells was significantly larger than that for the aggresome-free cells, the value increased dramatically for high-AgDD cells that formed aggresome spontaneously from peripheral aggregates without BTZ treatment (Fig. S1D). Therefore, we conclude that, although misfolded protein may enter the aggresome without the “pre-aggresome particle” stage under proteasome inhibition, efficient aggresome formation still requires a prior aggregation of the misfolded protein. It is not clear how proteasome inhibition loosens this requirement. One possibility is that misfolded AgDD may join the nonspecific protein aggregates induced by proteasome inhibition to be recognized by the transport machinery^35,36^.

To further study how the aggregation status of AgDD modulates its tendency to form aggresomes, we performed single-particle tracking of live-cell time-lapse images to determine the MTOC-directed velocity of individual AgDD aggregates at each frame (Fig. 1I). We then correlated the velocity with the aggregate diameter, which had been determined by fitting the aggregate’s image with a 2D-Gaussian function, upon calibration using fluorescent size standards (Fig. 1J, S3; Methods). Notably, the MTOC-directed velocity increased with the size of an aggregate, with a mean value of 0.6 nm/s for aggregates around 0.25 µm, rising to 1.4 nm/s for aggregates around 0.75 µm. Both values were significantly lower than the typical dynein velocity around 500 nm/s when transporting cellular organelles and vesicles of similar sizes (Fig. 1J)^37,38^. In summary, our results indicate that the sequestration of misfolded AgDD into the aggresome is likely to depend on the nucleation of AgDD into cytoplasmic aggregates. This dependence may be driven by the selectivity of the transport machinery, which favors larger aggregates.

### Reconstitution of dynein-mediated transport of protein aggregates in *Xenopus laevis* egg extract

Direct interpretation of the tracking result above is complicated by the potential involvement of the plus-end motor kinesins, changes in aggregate size, cytoplasm heterogeneity and other confounding factors from live cells. To investigate the mechanism of size selection in aggresome formation, we reconstituted the MTOC-directed transport of protein aggregates in *Xenopus laevis* egg extract (XE), an active cytoplasm that has been widely employed to study dynein activities in biological processes, such as spindle formation and the transport of organelles^37,39^.

Aggregation of purified AgDD in XE was triggered by adding recombinant FKBP(F36V) to deplete free Shield-1 (Supp. Video 2). AgDD aggregates were then diluted into interphase actin-depolymerized XE where the formation of microtubule asters was seeded by centrosomes from demembranated *Xenopus* sperms and probed with Alexa647-conjugated tubulin. The mobility of aggregates along microtubules was examined in a customized imaging chamber with dual-color fluorescence microscopy (Fig. 2A; Methods)^40^.

**Figure 2.**
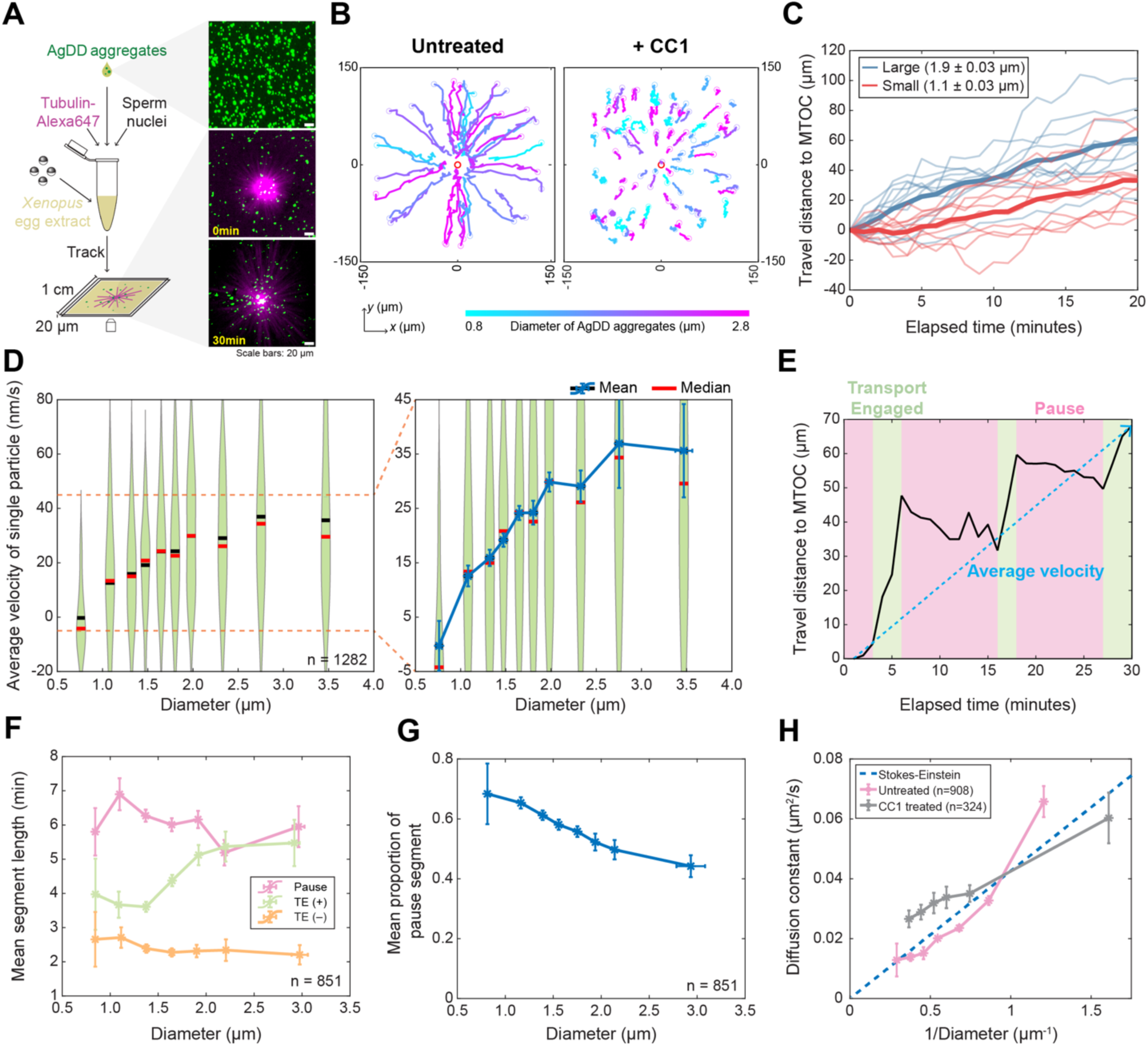
Reconstitution of dynein-dependent transport of AgDD aggregates in *Xenopus laevis* egg extract (XE) **(A)** Schematic and wide-field images of the aggregate transport assay. Recombinant AgDD (green) was induced to aggregate in XE and 1:20 diluted into a working XE containing Alexa647-labeled microtubule asters (magenta) and imaged in a 1 cm × 1 cm × 20 μm customized chamber once per minute for 30 minutes at 18 °C. **(B)** Example trajectories of AgDD aggregates. Measurement was performed as in **A**, with or without 40 μg/mL CC1 to inhibit dynein-dynactin in XE. Aggregates were randomly selected and colored by diameter. The MTOC was marked with an empty red circle at the center. Trajectories of 10 large and 10 small aggregates from the untreated group are plotted in **C** (mean diameter ± standard deviation in legend), with the averaged trajectories shown as thick lines. **(D)** The relationship between the time-averaged transport velocity and diameter of individual aggregates. The average velocity of individual aggregates was calculated as the total travel distance towards the MTOC divided by the trajectory’s duration (illustrated in **E**). The velocities of individual aggregates were grouped by the aggregates’ diameter in the violin plot. Right: a zoom-in plot showing the mean ± SEM within each group. A total of 1282 (n) aggregates around 16 microtubule asters from 2 batches of XE were included in the analysis. **(E)** An example trajectory of AgDD aggregate to illustrate the segmentation process. “Pause” and “Transport Engaged (TE)” segments were respectively colored in pink and green. **(F)** The relationship between the aggregate diameter and the lengths of the pause and TE segments, shown as mean ± SEM within each size group. 851 (n) trajectories from the experiment in **D** that can be tracked for longer than 15 minutes were selected and segmented as illustrated in **E**. TE (+): transport-engaged segment when the aggregate moves towards the MTOC; TE (-): transport-engaged segment when the aggregate moves away from the MTOC. **(G)** The relationship between the aggregate diameter and the proportion of the pause segment, using the same data from **F**. The plot shows the mean proportion ± SEM within each size group. **(H)** Aggregate’s diffusion constants during pause, with or without 40 μg/mL CC1 in XE. Indicated numbers (n) of aggregates that could be tracked for at least 15 minutes were selected and grouped by their diameters. Diffusion constants during pause were calculated as described in the methods and plotted against the inverse of the mean diameter of each size group. The dashed line represents the prediction by the Stokes-Einstein law as *k_B_T*/(3*πηd*), where *d* is the aggregate diameter, *η* is the dynamic viscosity of XE (0.01 Pa·S), *k_B_* is the Boltzman constant, and *T* is 291 Kelvin.

Upon incubation, we observed efficient MTOC-directed transport of AgDD aggregates resulting in most aggregates concentrated at the MTOC vicinity within 30 minutes, resembling aggresome formation in live cells (Fig. 2A, 2B; Supp. Video 3). Efficient transport was also observed in cycling XE with intact actin cytoskeleton (Supp. Video 4). This transport depended on a functional dynein-dynactin complex, as the addition of a p150 fragment CC1, widely used as a dynein inhibitor, blocked the transport, consistent with the previous observations with other aggregation-prone proteins in cells (Fig. 2B; Supp. Video 5). Processive backward (i.e., away from MTOC) transport of aggregates was not observed upon inhibiting dynein-dynactin, which suggests a lack of kinesin activity in the transport process, consistent with a general absence of kinesin activity in *Xenopus* eggs^37,41^. Interestingly, AgDD aggregates formed in a buffer solution showed no transport when incubated in XE (Supp. Video 6), suggesting that the cytoplasmic factors important for transport need to be present at the time of aggregate formation.

Although substrate ubiquitylation is important for aggresome formation by certain aggregation-prone proteins^3^, adding a high concentration of a nonspecific deubiquitylating enzyme Usp2^CD^ to deplete most Ub conjugates did not perturb the kinetics of AgDD aggregate transport in XE (Fig. S4A). In addition, we did not detect significant transport of Dynabeads conjugated with either K48- or K63-linked ubiquitin chains in XE (Fig. S4B). These results corroborate the findings in cells (Fig. 1C) to suggest a negligible role of ubiquitylation in AgDD aggresome formation.

### Average transport velocity of aggregates in *Xenopus* egg extract positively correlates with the size of aggregates

It took approximately 30 minutes for a peripheral aggregate to travel to the MTOC, much longer compared to the time constants of dynein molecule stepping or dynein-microtubule interaction, which typically last for seconds^42^. We therefore examined the transport phenomenon over both long- and short-time scales to identify the mechanisms contributing to size selectivity.

We first performed a long-term measurement at 1-minute per frame to record the entire transport process. We tracked the movement of individual AgDD aggregates, calculated the time average of each aggregate’s velocity towards the MTOC during tracking and associated the velocity with the aggregate diameter (Fig. 2C, 2D). Most aggregates’ size appeared stable during the experiment (Fig. S5) and fusion events were rarely detected, as the aggregate concentration was low. Like the transport of AgDD aggregates in cells, the average transport velocity in XE positively correlated with the aggregate size and approached zero when the aggregate diameter reduced to approximately 0.7 µm (Fig. 2C, 2D). In contrast, polystyrene beads, which non-specifically interact with cellular proteins, showed a negative size correlation in the MTOC-directed transport in XE (Fig. S6), suggesting that PSS in aggregate transport is mediated by specific interactions between dynein and the aggregates.

PSS was robustly observed across experimental replicates or using different batches of extracts (Fig. S7). Inhibiting the activities of the proteasome, Hsp70, or Hsp90 did not perturb size selectivity, whereas inhibiting the catalytic activity of HDAC6 reduced the average velocity and increased the size threshold for undergoing MTOC-directed transport, consistent with the role of HDAC6 in aggresome formation in cells (Fig. 1C, S8).

When traveling to the MTOC, AgDD aggregates exhibited “stop-and-go” movements (Fig. 2C, 2E). To determine which parameters of aggregate transport were modulated by size, we divided each trajectory into distinct kinetic segments. For segmentation, we fitted each trajectory with a piecewise linear function to achieve the normalized root mean square error (NRMSE) of 0.072 and designated a segment as “Pause” if the net displacement within this segment could be accounted by simple diffusion and measurement errors with a 95% confidence level (Fig. 2E; Methods). Otherwise, the segment was labeled as “Transport Engaged (TE),” to be differentiated from the processive movement at a finer timescale (see below). Aggregates alternated between these two states when moving towards the MTOC. The increase in aggregate size correlated with the lengthening of the TE segment, while the length of the pause segment remained stable (Fig. 2F). As a result, the mean proportion of the pause segment duration within a trajectory decreased with the aggregate size (Fig. 2G). These results reveal that larger aggregates tend to be more persistently engaged in MTOC-directed transport.

The pause segments are best characterized by a freely diffusing state, because the apparent diffusion constant of AgDD aggregates scaled inversely proportional to aggregates’ diameter during pauses, as predicted by the Stokes-Einstein equation, where the cytoplasmic coefficient of viscosity was taken as 0.01 Pa·s (Fig. 2H)^43^. Consistently, the diffusion constant in the presence of CC1, which inhibits dynein-dynactin, showed a similar dependence on aggregate diameter.

While aggregates in most TE segments moved towards the MTOC, brief backward movement was occasionally detected, accounting for an average of 8.6% ± 0.39% of the observation time (Fig. 2F, “TE (-)”). Kinesin-independent backward movement has been reported for the transport of dynein cargoes, although the underlying mechanism remains unclear^44–46^. We opted to neglect the contribution from backward segments in the following analysis, as they played a minor role in determining the average velocity and did not appear to vary with size (Fig. 2F). In summary, aggregate transport shows distinct features from the transport of conventional dynein cargoes, including a slow average velocity and a PSS. Both features can be recapitulated in the XE reconstitution system.

### The size-selection mechanism in protein aggregate transport

We next conducted high-resolution single-particle tracking experiments to investigate the detailed mechanism underlying the PSS in aggregate transport. AgDD aggregates were tracked at 30 frames per second for 2 minutes, with approximately 10 nm uncertainty in sub-pixel localization as determined using surface-immobilized fluorescent beads (see Methods). These experiments confirmed the positive correlation between average transport velocity and aggregate size (Fig. 3A, 3B) and uncovered additional kinetic features that are important for understanding the size-selection mechanism.

**Figure 3.**
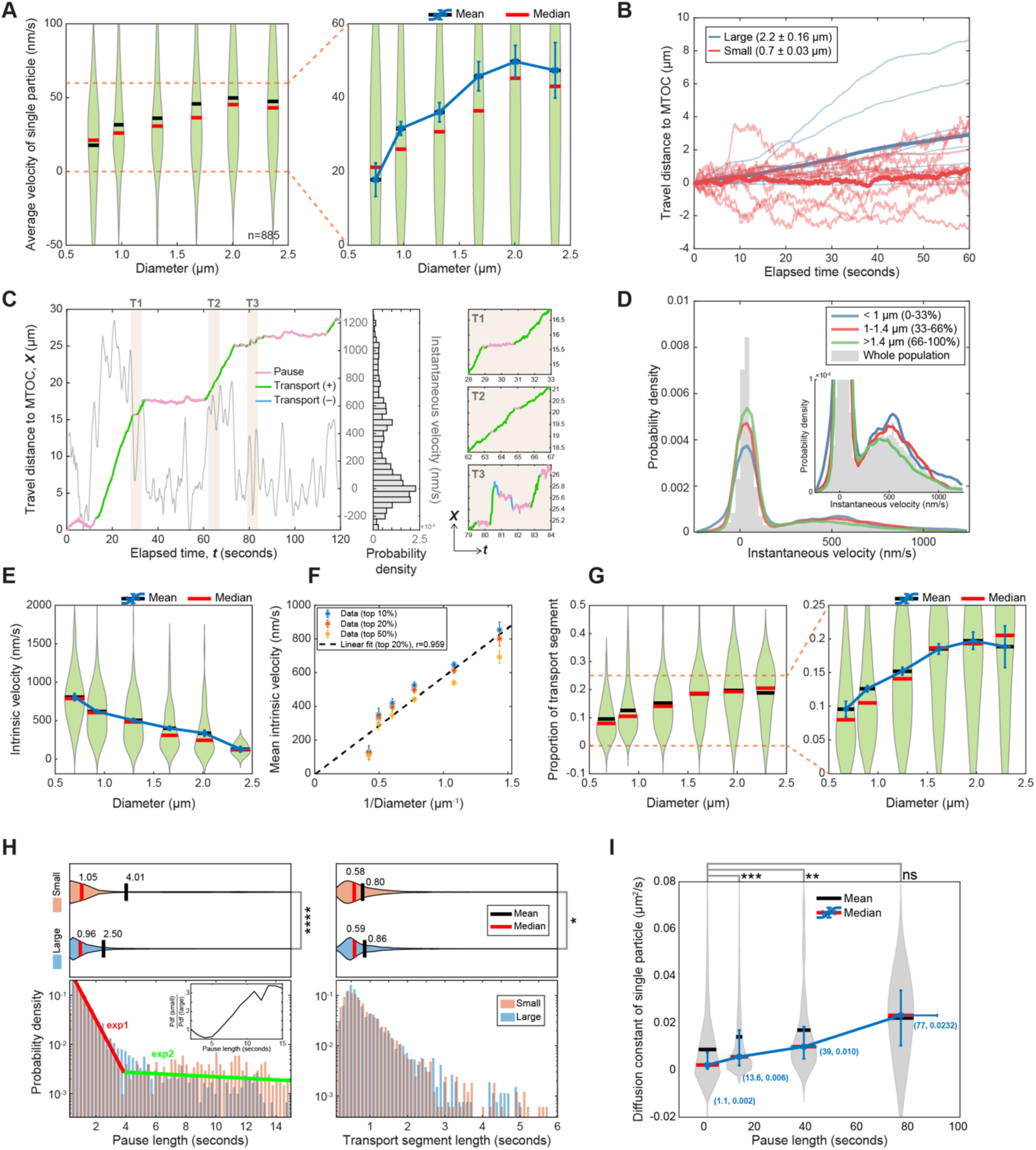
Positive size selectivity in aggregate transport originates from a size-dependent increase in the likelihood of engaging in active transport. **(A)** The relationship between the time-averaged velocity and the diameter of individual aggregates, presented in a violin plot with a zoom-in axis on the right. 885 trajectories around 6 microtubule asters were acquired at 30 frames per second for 120 seconds. Error bars represent the SEM within each size group. **(B)** Example trajectories of AgDD aggregates. Data were from the experiment in **A**. 10 large aggregates (blue) and 10 small aggregates (red) were randomly chosen and plotted as faint lines (mean diameter ± standard deviation in legend). The averaged trajectories of the large and the small groups were shown as thick lines. **(C)** An example trajectory of AgDD aggregate to illustrate the segmentation procedure. “Pause”: pink; “Transport (+)”: moving towards the MTOC, green; “Transport (-)”: moving away from the MTOC, blue. The instantaneous velocity was calculated as the time derivative using a 2-s rolling time window, overlaid (gray curve) on the right axis, and the probability density is shown on the side. Zoom-in windows of the shaded regions, marked by T1, T2 and T3, are shown on the right. **(D)** The probability densities of instantaneous velocities of AgDD aggregates calculated as in **C** using only the “Transport (+)” segments. 885 trajectories from **A** were divided into three groups by aggregates’ diameter: < 1 μm (0-33 percentile); 1-1.4 μm (33-66 percentile); > 1.4 μm (66-100 percentile). The probability density of the entire population is overlaid as gray bars. Inset: a view with a local *y*-axis. **(E)** The relationship between the intrinsic velocity and the diameter of AgDD aggregates, presented in a violin plot. Intrinsic velocity was defined for each aggregate as the mean of the top 20% of instantaneous velocities calculated as in **D**. The mean intrinsic velocity within each size group was plotted against the inverse of the aggregate diameter in **F**. Similar results were obtained using the top 10% and top 50% of instantaneous velocities in the definition of intrinsic velocity. Error bars represent the SEM in each group. Data were from the experiment in **A**. **(G)** The relationship between the proportion of the transport segment and the diameter of AgDD aggregates in a violin plot. Trajectories in **A** were grouped by size and segmented into transport (+/-) and pause segments as illustrated in **C** (Methods). The mean proportion within each size group was highlighted on the right. Error bars represent the SEM in each group. **(H)** Probability densities of the lengths of pause and transport segments. Trajectories of the largest 10% and the smallest 10% aggregates in **A** were included in the analysis. The mean and median values of the segment lengths were labeled on the top panel. Lower panel: time constants for the large aggregates (0.77 ± 0.03 s and 7.24 ± 3.12 s at the 95% confidence level) and the small aggregates (0.76 ± 0.04 s and 27.79 ± 14 s) were from fitting their pause length distributions with a double exponential function. The red and blue lines represent the two exponential modes of the large aggregates in a semi-log plot. Inset: ratio of the pause length distribution of the small aggregates over that of the large aggregates. The statistical significance of the difference between groups of large and small aggregates was calculated using Student’s two-tailed *t*-test (\**p* = 0.036; \*\*\*\**p* < 0.0001). **(I)** The relationship between the diffusion constants of AgDD aggregates during pause and the pause length. The apparent diffusion constant during each pause segment was calculated based on a linear regression of the mean square displacement on time, grouped by the pause length, presented in a violin plot. For each group, the median values of the pause length and diffusion constant were labeled next to the distribution and the error bars represent the first and third quartiles. The statistical significance of the difference between groups was calculated using Student’s one-tailed *t*-test (ns: *p* = 0.16; \*\**p* = 0.01; \*\*\**p* < 0.001). Data were from the experiment in **A**.

Larger aggregates may be carried by a greater number of dynein molecules, leading to faster transport. Alternatively, their size could predispose them to remain longer in the transport mode. To test the former possibility, we analyzed the instantaneous velocity calculated as the time derivative of an aggregate’s trajectory (Fig. 3C). Although the time-averaged velocity of an aggregate was low, the instantaneous velocity spread significantly higher, approaching the typical dynein transport velocity around 500 nm/s (Fig. 3C, 3D). To circumvent the influence of pausing during transport and stochasticity, we chose the mean value of the top 20% instantaneous velocities within transport segments in a trajectory to represent the velocity of that aggregate after forming an active transport complex with dynein and microtubule and called it “intrinsic velocity.” Remarkably, the intrinsic velocity correlated inversely with aggregate size, aligning with the expectation by the viscosity rule (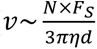, where *N* is the number of dynein; *F*_*S*_ is the dynein stalling force; *η* is the viscous coefficient; *d* is the diameter) (Fig. 3E, 3F)^25,38^. Choosing different instantaneous velocity thresholds to define the intrinsic velocity led to the same conclusion (Fig. 3F). This observation also indicates that the average number of dynein molecules actively hauling an aggregate does not change significantly with its size.

To understand how a PSS in the average velocity emerges from a NSS in the intrinsic velocity, we delved deeper into the transport kinetics. Notably, under high temporal resolution, the movement of most aggregates appeared highly episodic. Trajectories with a net MTOC-directed movement over a 2-minute observation period, corresponding to the TE segments defined previously (Fig. 2E), are indeed composed of numerous short, processive transport segments interspersed with pauses of varying lengths (Fig. 3B). The shape of the trajectory also varied with the aggregate size, with larger aggregates’ trajectories appearing “smoother” (Fig. 3B). We applied a piecewise linear function with up to 153 segments to fit each trajectory, achieving a NRMSE of 0.0217. A segment was classified as “Transport,” if it exhibited a nonzero transport velocity with 90% confidence (Fig. 3C; Methods); otherwise, it was classified as “Pause.” By analyzing the proportion of each kinetic mode, we concluded that larger aggregates were more likely to reside in the transport state (Fig. 3G). This gain of transport probability was sufficiently large to offset the decrease in intrinsic velocity of larger aggregates due to viscosity.

We next examined the time distribution of different kinetic segments to understand their contributions to the size-dependent modulation of the transport likelihood. The distribution of pause segment length was best described by a double exponential function, containing distinctly separated time constants. The fast time constant was 0.77 ± 0.03 seconds and remained invariant across sizes, whereas the slow time constant was about 3.8 times longer for the smallest 10% of the aggregates compared to the largest 10%, resulting in a significantly shorter mean pause length for the larger aggregates (Fig. 3H). These findings indicate that larger aggregates more readily resume transport after pause. The distribution of transport time did not appear to vary with size, as the smallest 10% of aggregates had an essentially identical distribution as the largest 10% (Fig. 3H).

In addition, we found that the apparent diffusion constant of aggregates in the pause state appeared to correlate with the pause duration: shorter pause segments were predominantly associated with slower diffusion, suggesting a stationary or motion-restricted state (Fig. 3I) (see Discussion). In summary, a diffusing aggregate engages with the transport machinery for multiple rounds *en route* to the MTOC, and its movement is episodic at both long and short timescales. The PSS in transport velocity is primarily caused by the greater propensity of larger aggregates to participate in active transport, which effectively offsets the reduction in their intrinsic velocity due to viscosity.

### Aggresome adaptors mediate episodic transport in *Xenopus laevis* egg extract

The size-dependent modulation of the transport likelihood requires the likelihood value to be non-saturating or significantly less than one in the typical size range, which can be achieved through transport episodicity. We next studied whether transport episodicity depends on the properties of cargo or cargo adaptors. To this end, we used a chemically defined cargo with specific adaptors.

We first identified factors mediating the aggregate transport in XE through characterizing the interactome of AgDD aggregates. We aggregated AgDD in XE and coupled the aggregates to Dynabeads, using soluble AgDD that was not transported by dynein as a control. After incubation in XE, the bead-bound proteins were analyzed by Tandem-Mass-Tag mass spectrometry (TMT-MS). This analysis identified several protein chaperones that were enriched with AgDD aggregates, including Hsp90, DNAJAs, and the known aggresome mediators Hsp70 and 14-3-3, together with dynein-dynactin subunits (Fig. S9; Table S1; Supp. Data File 1). Consistent with DD-degron misfolding, proteasome subunits were also identified.

Several chaperonin (TRiC/CCT) subunits also showed enrichment with AgDD aggregates. Previous studies discovered that chaperonin subunits were recruited to the aggresome and interacted with DIC, suggesting a potential role of the chaperonin in aggresome formation^10,48,49^. We found that the chaperonin subunits, CCT2 and CCT8, colocalized with the aggresome formed by poly-Q protein Htt (Q94)-CFP in HEK293T cells using immunofluorescence (Fig. S10G), though marked colocalization between chaperonin subunits and AgDD aggresomes was not observed. To further explore chaperonin’s role in aggresome formation, we transiently knocked down CCT2, CCT4, or CCT8 in HEK293T cells by RNAi and studied the consequence on aggresome formation (Fig. S10A, S10B). Depleting chaperonin subunits resulted in a modest, yet statistically significant, delay in AgDD aggresome formation, with little impact on cell growth (Fig. S10C, S10D). Similarly, a minor delay was observed in cells depleted of HDAC6 or BAG3 (Fig. S10B, S10D). Consistent with the involvement of redundant aggresome pathways, double knockdowns of aggresome adaptors resulted in longer delays in aggresome formation (Fig. S10B, S10D). This delay is not unique to the AgDD system, as chaperonin knockdown also retarded poly-Q aggresome formation in HEK293T cells (Fig. S10E, S10F).

We then used beads coated with the identified factors or known dynein adaptors, to address whether these factors can mediate cargo transport in XE and whether they incur different transport kinetics which may be associated with distinct size selectivity. We prepared 2.8 µm-diameter fluorescent beads coated with specific antibodies to precipitate different factors from HeLa cell extract (Fig. 4A). TMT-MS with absolute protein level quantification verified that each antibody-coated bead primarily precipitated its designated antigen (Table S2; Supp. Data File 2). No dynein or dynactin component was detected on beads by TMT-MS, suggesting their low abundance.

**Figure 4.**
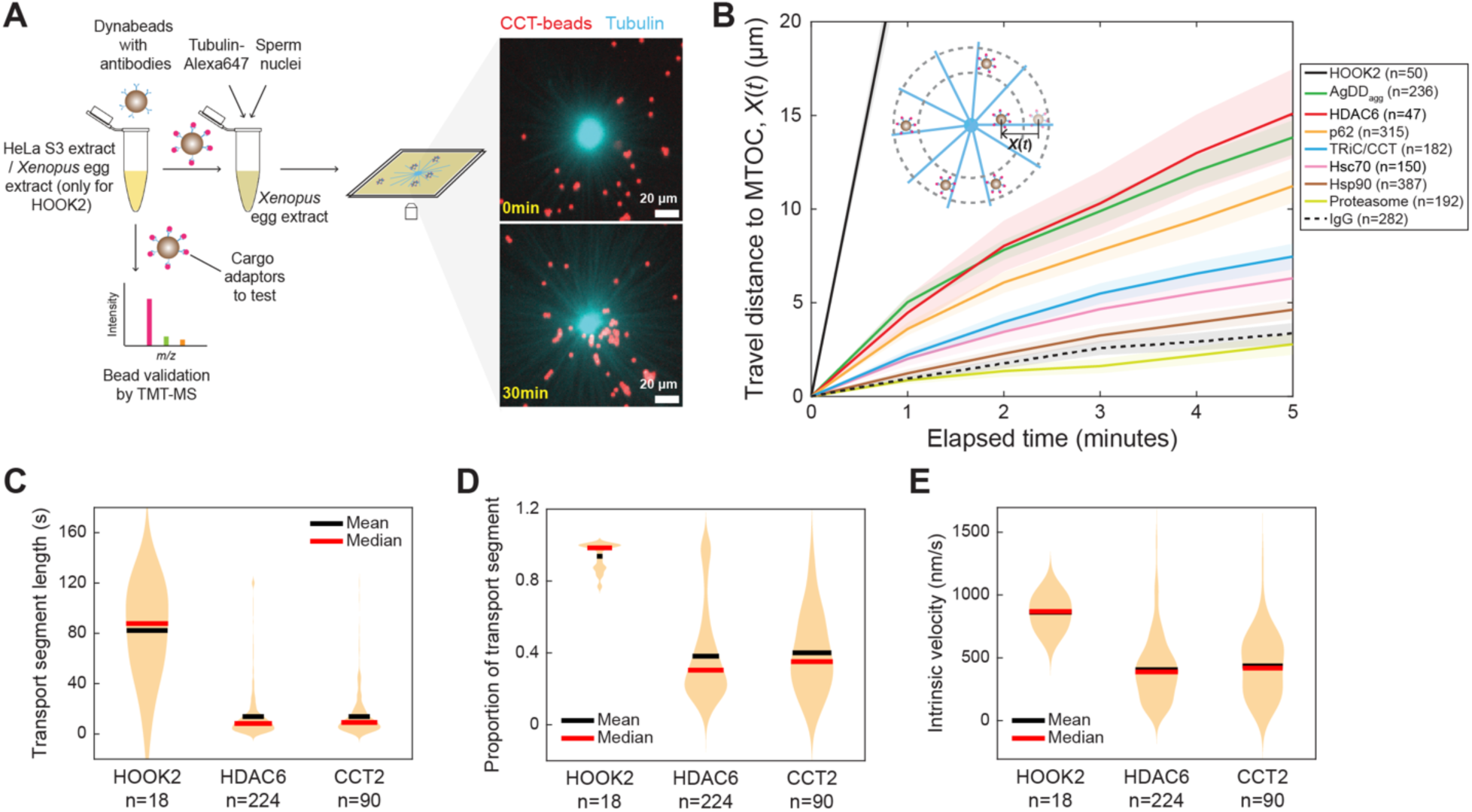
Aggresome adaptors mediate dynein-dependent transport in *Xenopus laevis* egg extract (XE) **(A)** A schematic workflow of studying the transport of adaptor-coated beads in XE. Adaptor-coated beads were incubated in interphase XE with Alexa647-labeled microtubule asters (cyan), imaged every 1 minute for 30 minutes at 18 ℃. Example images from an experiment with chaperonin (CCT) coated beads (red) are shown on the right. **(B)** The averaged trajectories of different factor-coated beads in XE. Experiments were performed as described in **A**. Transport of AgDD aggregates (AgDD_agg_) was included for comparison. Line shading represents SEM. n: number of trajectories. **(C)** Transport segment length distribution for beads coated with indicated factors in a violin plot. Experiments were performed as in **A**, but acquired at 30 frames per second for 120 seconds. The distributions of the transport segment proportion and the intrinsic velocity were presented in **D** and **E** respectively. Intrinsic velocity was defined as the mean of the top 20% of instantaneous velocities as in Fig. 3E. n: number of trajectories.

Upon incubating adaptor-coated beads in XE containing microtubule asters, we recorded efficient MTOC-directed transport of beads coated with HDAC6, SQSTM1/p62, and Hsc70 at an average velocity of 21∼50 nm/s, similar to the transport of AgDD aggregates at 11∼32 nm/s (Fig. 4B). These activities of known aggresome adaptors support the biological relevance of our reconstitution system. Notably, we recorded efficient MTOC-directed transport of CCT2-coated beads at a similar velocity (Fig. 4A, 4B; Supp. Video 7). These results suggest a likely role of the chaperonin, or its subcomplex, as another dynein adaptor for protein aggregates. In contrast, the transport of proteasome- or Hsp90-coated beads was inefficient and indistinguishable from that of random-IgG-coated beads (Fig. 4B), suggesting these factors may not directly engage with the transport machinery.

Beads coated with the dynein-activating adaptor HOOK2 traveled at 480 nm/s, much faster than beads coated with the aggresome adaptors (Fig. 4B). Consistently, at high temporal resolution, the movement of aggresome adaptor-coated beads was highly episodic, characterized by short processive runs punctuated by frequent pauses, akin to the transport of AgDD aggregates (Fig. 4C). In contrast, HOOK2-coated beads exhibited a higher intrinsic velocity, and the movement was mostly devoid of short pauses (Fig. 4C, 4D, 4E). Overall, the transport mode accounted for approximately 40% of the observation time for the aggresome adaptor-coated beads, whereas HOOK2-coated beads consistently maintained their transport velocity (Fig. 4D). The similarity in transport kinetics of AgDD aggregates and aggresome-adaptor-coated beads suggests that the aggregate transport in XE is likely to be mediated by the aggresome adaptors.

In conclusion, our results show that aggresome adaptors mediate episodic transport of protein aggregates, resulting in a lower likelihood of the cargo staying in the active transport state.

### Dynein’s activating adaptor perturbs the normal size selectivity in aggresome formation

Since the PSS is mainly driven by size-dependent modulation of the transport proportion (Fig. 3G), we reason that the episodic kinetics by aggresome adaptors could be the key to the PSS in aggregate transport. To examine whether the PSS in aggregate transport is specifically associated with the aggresome adaptors, we tethered dynein’s activating adaptors, HOOK2 or HOOK3, to protein aggregates by expressing AgDD fused with these adaptors in U2OS cells and studied the transport of the corresponding substrates. Anti-GFP immunoblotting suggested that most GFP signal was from the full-length fusion protein (Fig. 5A).

**Figure 5.**
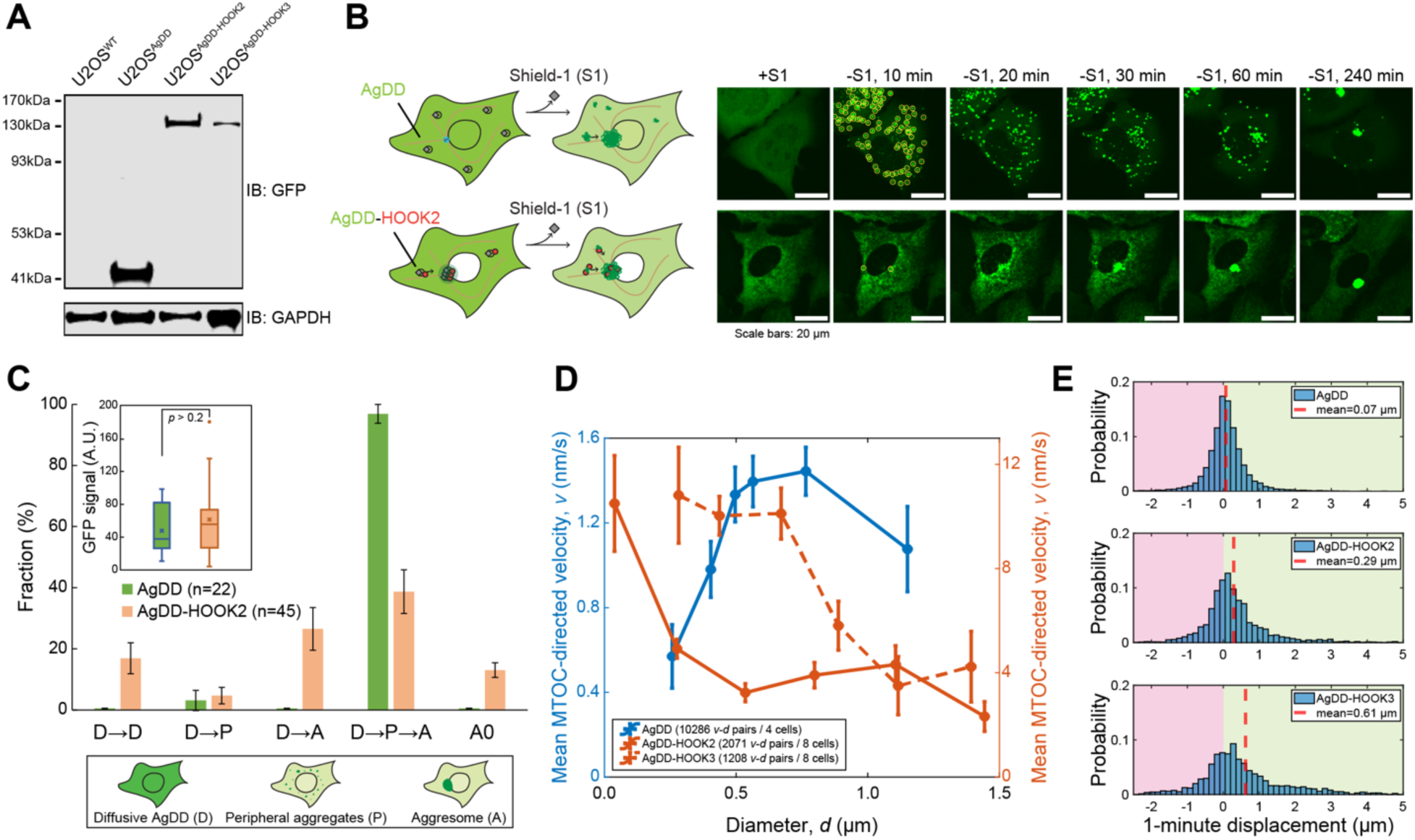
Dynein’s activating adaptors HOOK2 and HOOK3 are associated with a negative size selectivity in aggregate transport. **(A)** Anti-GFP western blot of U2OS cells stably expressing AgDD (41 kDa), AgDD-HOOK2 (124.5 kDa) and AgDD-HOOK3 (124.4 kDa). Blot was reprobed with anti-GAPDH as the loading control. **(B)** Schematic and live-cell confocal images of U2OS cells stably expressing AgDD (top) or AgDD-HOOK2 (bottom), before and after Shield-1 removal. Peripheral aggregates were identified by *TrackMate* and circled in yellow at “10 min” as an example where 55 particles were identified in the AgDD cell, and 2 in the AgDD-HOOK2 cell. **(C)** Classification of the aggregation-to-aggresome phenotypes, presented as in Fig. 1F. Bar plot shows the mean ± SEM, over 4 different FOVs. n: number of cells analyzed. D, P, A are defined as in Fig. 1F. A0: perinuclear puncta detected in the presence of Shield-1. Inset: cellular GFP signal in the analyzed AgDD and AgDD-HOOK2 cells before Shield-1 removal. P-value > 0.2 by unpaired Student’s two-tailed *t*-test. Boxplot shows mean (“x”), median (-), first and third quartiles; upper/lower whiskers extend to 1.5 × the interquartile range. **(D)** The relationship between the mean MTOC-directed velocity and the aggregate diameter, as presented in Fig. 1J. Error bars represent the SEM within each size group. **(E)** Probability distributions of the MTOC-directed displacement over 1 minute of AgDD, AgDD-HOOK2, and AgDD-HOOK3 aggregates in the experiment in **D**. Dashed red lines mark the mean values.

Upon Shield-1 removal, AgDD-HOOK2 rapidly accumulated at the perinuclear site that resembled an aggresome (Fig. 4B; Supp. Video 8). We identified the aggregates in each frame using the ImageJ plugin “*Trackmate*,” and found that, in 40.7% ± 4.62% of cells with an aggresome, fewer than 3 peripheral aggregates were detected before the appearance of the perinuclear punctum (Fig.5C, group “D→A”), while a typical AgDD cell showed more than 40 peripheral aggregates in this process. A perinuclear punctum of AgDD-HOOK2 was detected even in the presence of Shield-1 in 13.1% ± 1.36% of cells, much more frequently than in AgDD cells with a similar GFP level (Fig. 5C, group “A0”; Supp. Video 8). These distinct features of the aggresome formation from AgDD-HOOK2 indicate that diffusive AgDD-HOOK2 or small aggregates that are below our detection limit can be sequestered into the aggresome in these cells.

In cells that formed distinguishable peripheral AgDD-HOOK2 aggregates, we tracked aggregate movement and recorded a significantly higher transport velocity compared to the AgDD aggregates (Fig. 5D, 5E). Remarkably, AgDD-HOOK2 aggregates exhibited a NSS, with larger aggregates being transported more slowly, akin to the transport of conventional dynein cargoes (Fig. 5D, right axis). AgDD fused with another activating adaptor HOOK3 had a similar phenotype. Therefore, we conclude that the PSS during aggregate transport is uniquely associated with the aggresome adaptors, as coupling activating adaptors to protein aggregates perturbs the normal substrate selectivity in aggresome sequestration.

### Physical modeling suggests that the stability of the cargo-dynein-microtubule complex determines the size selectivity in cargo transport

Although size-dependent phenomena are universal in biological systems, few biological mechanisms have been identified to account for size selectivity. We speculate the observed PSS has a physical origin and asked whether we could establish a simple physical model to account for our main observations. To engage in active transport, a freely diffusing protein aggregate must first be captured by a dynein motor and loaded onto the microtubule. Most aggregates switch between the previously defined transport-engaged (TE) state and the freely diffusing state multiple times before reaching the MTOC. Movement in the TE state is not free of interruption but is instead characterized by frequent short pauses, which can be caused by the dissociation of dynein from either the cargo or the microtubule. Rapid reformation of the active transport complex consisting of cargo, dynein, and microtubule after a short pause indicates that the aggregate may remain in the vicinity of microtubules during this process, which is also supported by the slow diffusion observed during the short pause (Fig. 3I)^50^. We therefore call the above process a “Local Dynein Cycle” (Fig. 6A). The resulting episodic transport enables size-dependent modulation of the likelihood of the cargo dwelling in the transport mode, which is the primary mechanism for the PSS in aggresome formation. Larger aggregates, despite slower diffusion, may be more efficiently captured by microtubules (or by microtubule-dynein). Their larger surface area and potentially more bound aggresome adaptors could also favor dynein’s local reattachment. This size-dependent multivalency effect is akin to the avidity-entropy mechanism that enhances a ligand’s effective affinity with multivalent receptors^51,52^.

**Figure 6.**
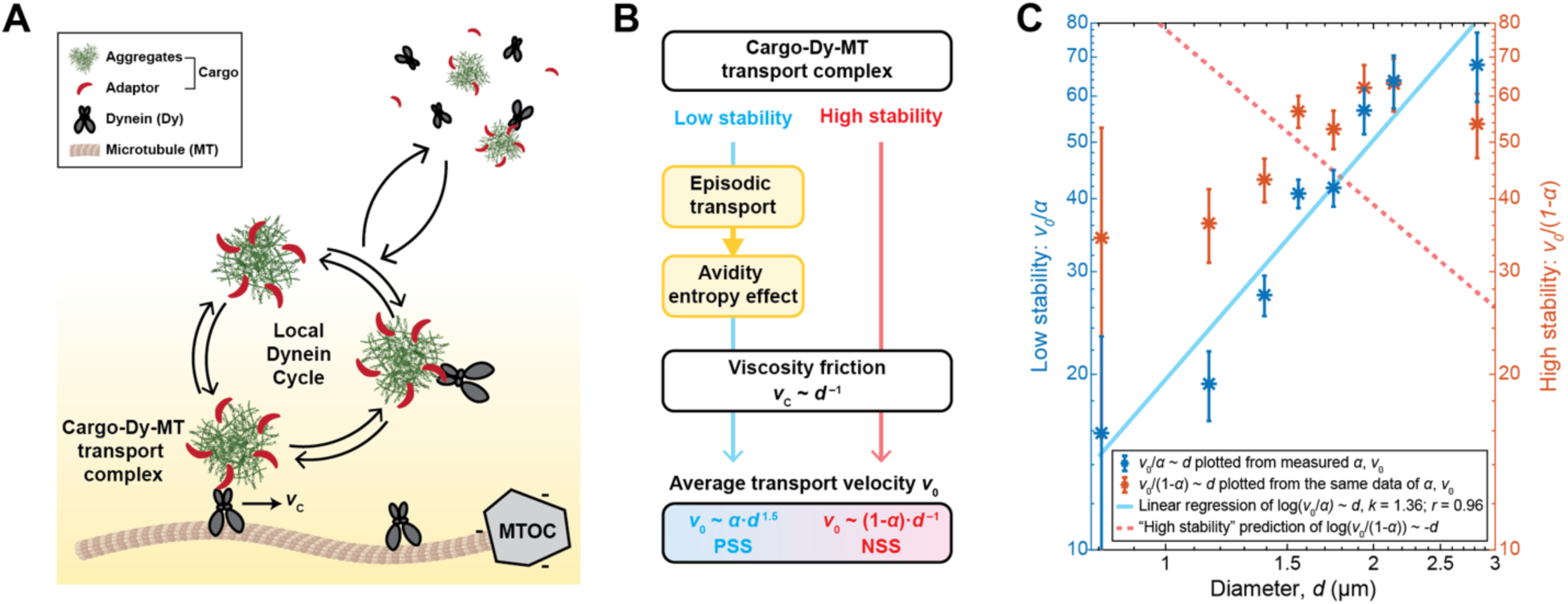
A physical model of dynein-mediated transport suggests the stability of the active transport complex determines cargo size selectivity. **(A)** Schematic of the dynein transport model. See text for details. The transport complex formed by cargo, dynein, and microtubule is labeled on the graph. *v*_C_: intrinsic velocity of dynein transport. **(B)** Diagram showing the mechanisms that collectively determine the size selectivity in cargo transport when the transport complex has low or high stability. The scaling expression of the average transport velocity *v*_0_ as a function of cargo diameter *d* was derived under two scenarios: 1) When the transport complex is transient or has low stability, *v*_0_ varies in proportion to *α* · *d*^1.5^, which gives rise to PSS. *α* is the probability of the freely-diffusing state as a function of *d* (Fig. 2G); 2) When the transport complex has high stability, *v*_0_ varies in proportion to (1 − *α*) · *d*^-1^, which gives rise to NSS. See Methods for details. **(C)** Comparison of the model predictions with experimental measurements. *v*_0_and *α* were obtained from Fig. 2D**, 2G**. Experimental results are replotted as the mean ± SEM of either *v*_0_ ∕ *α* (blue, left axis, by the low-stability expression) or *v*_0_ ∕ (1 − *α*) (red, right axis, by the high-stability expression), against *d* in a log-log plot. Blue line represents the linear regression using the low-stability expression with the fitted slope 1.36 and *r* value 0.96. The red dashed line is the best fitting result by the high-stability expression (slope = -1).

We next address the important question of whether this conceptual model is physically achievable and whether it is sufficient to account for the PSS in aggregate transport. Specifically, can the size-dependent modulation of transport likelihood through dynein rebinding cycles provide enough size bias to offset the reduction in intrinsic velocity due to viscosity?

Furthermore, which system parameter dictates different size selectivities? To address these questions, we rewrote the conceptual model in Fig. 6A using statistical physics, which predicted how an aggregate’s average transport velocity should vary with its size (see Methods).

In this exercise, we recognized that the stability of the cargo-adaptor-dynein-microtubule transport complex critically determined the size selectivity and derived the analytic expressions for the average velocity *v*_0_under two conditions (Fig. 6B): 1) Transient transport complex, whereby most cargoes are not engaged in active transport under a steady state. We obtained 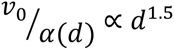, i.e., a PSS, where *d* is the diameter of the aggregate and *α*(*d*) is the likelihood of cargo dwelling in the freely diffusing pause state (Fig. 2G); 2) Stable transport complex, whereby most cargoes in the local dynein cycle are engaged in active transport. This gave rise to 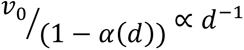, i.e., a NSS if *α*(*d*) does not change dramatically with the cargo size, as we observed for AgDD aggregates. It is worth noting that these scaling exponents are determined by the system’s geometry rather than kinetic parameters in the model. The experimentally determined exponent of 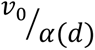 on *d* is 1.36 (Fig. 6C), closely aligning with the theoretical prediction for the first scenario and indicating a low stability of the transport complex. Low transport complex stability is also corroborated by the observation that aggregates and aggresome-adaptor-coated beads were transported in short bursts. In contrast, the movement of HOOK2-coated beads was more processive with fewer short pauses, suggesting a more stable transport complex. The stability of the active transport complex potentially accounts for the NSS in the transport of AgDD-HOOK2/HOOK3 aggregates.

## Discussion

Efficient targeting of misfolded proteins into the appropriate quality-control pathway is crucial for maintaining protein homeostasis. Aggresome formation preferentially sequesters aggregated misfolded proteins over their soluble counterparts. To elucidate the mechanism underlying the “size filter” effect, we reconstituted the dynein-mediated MTOC-directed transport of protein aggregates in a cell-free system, gaining valuable mechanistic insights.

Transport of protein aggregates by dynein exhibits exquisite features, highlighting a PSS, in contrast to the transport of conventional dynein cargoes where transport velocity and cargo size negatively correlate. The PSS in aggregate transport appears to be specifically associated with the aggresome adaptors as artificially attaching an activating adaptor to the aggregates reversed size selectivity. In searching for the size selection mechanism, we found that a simple physical model was able to recapitulate the scaling of the average transport velocity with aggregate size. This model suggests that the PSS in aggregate transport may not be attributed to a specific aggresome adaptor. Rather, it is a general feature originating from the recurrent formation of the cargo-dynein-microtubule transport complex. This rebinding process can manifest an avidity entropy effect that favors larger aggregates in the transport state to overcome the reduction in their intrinsic velocity. Similar mechanisms have been proposed to enhance the receptor-ligand and enzyme-substrate interaction through multivalency^51–53^.

To allow size-dependent modulation, the likelihood of cargo dwelling in the transport mode should not saturate, to maximize the avidity entropy effect. Having a low likelihood of transport, as mediated by the aggresome adaptors, requires the cargo-dynein-microtubule transport complex to be transiently formed or unstable. Although he molecular basis for this condition has not been clearly elucidated, HDAC6 has been suggested to interact weakly with dynein^13^. Such weak dynein interaction may be common among the aggresome adaptors, since they do not share the coiled-coil domain in the dynein-activating adaptors which extensively interacts with dynein-dynactin. The interaction between the aggresome adaptor and aggregates may also contribute to the instability of the transport complex. For example, previous studies suggest that HDAC6 interacts with the free C-terminus of Ub (or Ub chains) trapped in aggregates^22,54^. The high concentration of free Ub in the cytosol is likely to compete in this process, effectively weakening the interaction between HDAC6 and aggregates. In addition, a dynein motor associated with an aggresome adaptor may have lower processivity on the microtubule, compared to dynein complexed with an activating adaptor^19,55^. Interestingly, dynein appears to transiently interact with certain endosomal cargoes in HeLa cells and the interaction is presumably mediated by the activating adaptors^50^. The stability of the transport complex may also be modulated by dynein cofactors or through post-translational modifications^47^. These effects may collectively strengthen the avidity entropy effect.

Maximizing the avidity entropy effect through a transient transport complex also requires the cargo to remain close to microtubules after the transport complex disassembles, to allow efficient transport complex reformation. This requirement may be achieved through different mechanisms. A super-resolution microscopy study observed that endosomal particles in transit remain close to microtubules after dynein dissociation in live cells^50^. This dynein-independent microtubule tethering may be mediated by dynactin subunit p150^56^. Alternatively, cargoes may dwell close to the microtubule simply due to confined diffusion^57^. The microtubule surface also appears to engage beads coated with a variety of proteins to perform one-dimensional diffusion along the microtubule^58^. These mechanisms may prolong the local dynein cycle and enhance the size selectivity in transporting aggregates.

In the current model, we only considered transport mediated by a single dynein molecule. The inverse correlation between the intrinsic velocity and aggregate size suggests that the number of dynein molecules mediating transport does not change significantly with size. The occasional processive backward transport was also consistent with the behavior of single dynein reported in previous studies^44^. We cannot exclude the possibility of multi-dynein-mediated transport, especially when the dynein density is high on the cargo surface. Another limitation of the model is that it does not take into account any sudden change in the behaviors of dynein while remaining bound. Such an effect may not be directly involved in cargo size selectivity.

Protein aggregates are an important class of dynein cargo. Besides selectively targeting misfolded proteins into the aggresome, there may be other important advantages associated with the PSS. We previously reported that small clusters of aggregation-prone proteins are permanently present in the cell and serve to nucleate aggregates upon proteotoxic conditions^35^. The PSS may be necessary to retain these nucleation sites throughout the cytoplasm to capture abnormal polypeptide species in the emerging stress. Our study indicates that the choice of cargo adaptor is constrained by a tradeoff between the transport speed and size selectivity: activating adaptors may allow faster transport but favor smaller cargos; on the contrary, when PSS is desired, an aggresome adaptor can be employed, but at the expense of the transport velocity.

Multiple aggresome adaptors participate in transporting protein aggregates while their substrate specificities are not fully understood. In this study, we identified that the chaperonin complex is sufficient for mediating cargo transport by dynein and contributes to efficient aggresome formation in cells. In addition to PQC, many biological processes also involve the transport of protein aggregates or condensates. The avidity entropy effect we identified may be universal and can be applied to understand the specificity of other biological pathways processing protein aggregates.

## Data availability

Stable cell lines and plasmids generated in this study are available from the corresponding authors with a completed material transfer agreement. All other data supporting the findings of this study are available from the corresponding authors on reasonable request. Source data are provided with this paper.

## Code availability

All MATLAB codes for microscopy data analysis and data presentation are available from the corresponding authors upon request.

## Acknowledgements

We are grateful for funding from the National Institute of General Medical Sciences (R01 GM134064-01 to YL; R35 GM131753 to TM) and want to thank Lisa McCalla for administrative support. We are grateful for Westlake Fellowship to RF. We want to thank Daniel Finley, Tinting Yao, Louis Colson, Randall King, Mark Verheijen, Donghoon Lee, Thuan Beng Saw, Shutao Qi, and Shih-Han Lin for commenting on the manuscript. We want to thank the Core for Imaging Technology & Education (CITE) at Harvard Medical School for assisting with live-cell imaging and thank the staff at the Westlake University Microscopy Core Facility for advice and assistance in light microscopy data collection. We also thank Rachael Jonas-Closs for excellent *Xenopus* husbandry.

## Author contributions

R. F. and Y. L. conceived and designed the project. R. F., L. B., B. L., M. Z., and Y. C. carried out the experiments. K. D. and J. P. processed mass spec samples and data analysis. All authors participated in manuscript preparation.

