## Supplementary material for "Episodic Transport of Protein Aggregates Achieves a Positive Size Selectivity in Aggresome Formation": Materials and Methods

### Plasmids

AgDD-sfGFP was cloned from the original plasmid (Addgene plasmid #78289) into the pLenti6.3/V5-DEST gateway vector (Invitrogen) under the control of a CMV promoter. An HA-tag was inserted at its N-terminus. The open reading frames of human HOOK3 (Thermo Ultimate™ Human ORF Clones, #HORF96) and human HOOK2 (WZ Biosciences Inc.) were amplified and appended to the C-terminus of AgDD-sfGFP to create AgDD-sfGFP-HOOK3 and AgDD-sfGFP-HOOK2. HaloTag-Histone H2A in a retroviral vector with a hygromycin selection marker was from the Mitchison lab[1]. Htt (Q94)-CFP in a lentiviral vector under the control of a TetOn promoter was reported in a previous study[2]. The construct for purifying recombinant cys-$Ub^{L73F}$ was described previously[2]. The construct for expressing recombinant HA-AgDD-sfGFP in *E. coli* was made by inserting the target gene into the pTXB1 vector (NEB) through the SapI, NdeI restriction sites. Histone H2B-mCherry in the CSII-EF lentiviral vector was a gift from Tobias Meyer[3]. 6xHis-FKBP(F36V) in the pET15b vector was a gift from Thomas Wandless (Addgene plasmid #73180; http://n2t.net/addgene:73180; RRID:Addgene_73180)[4].

### Recombinant Protein Purification

6xHis-FKBP(F36V) was purified from *E. coli* cells using His60 Ni Superflow Resin (Takara Bio, #635662) according to the manufacturer's instructions. Eluted proteins were buffer-exchanged and concentrated to 1.75 mM in PBS + 10% glycerol and stored at -80 °C for future use. Protein concentration was determined using the Bio-Rad protein assay.

HA-AgDD-sfGFP was purified from *E. coli* cells via the intein-tag at the C-terminus on Chitin Resin (NEB, #S6651L). 20 mL of resin was used for 1 L of cell culture. Cells were lysed by sonication in the wash buffer (30 mM Tris-HCl pH 8.0, 300 mM NaCl, 2 mM $MgCl_2$, 5% glycerol, 1 mM PMSF, 0.5 mM EDTA) plus 0.5% Tween-20 and centrifuged at 35,000×g for 30 minutes at 4 °C. The supernatant was incubated with the Chitin beads (NEB, #S6651L) for 1 hour at 4 °C. Beads were washed twice with the wash buffer + 0.5% Tween-20 and twice with the wash buffer before being resuspended in the cleavage buffer (wash buffer + 50 mM dithiothreitol (DTT), 20 μM Shield-1) of 3 times the bead volume and rotated at 4 °C for 72 hours to release HA-AgDD-sfGFP. Flow-through was collected and proteins were further eluted three times using the wash buffer (+ 20 μM Shield-1) of 1/3 of the bead volume, with 10-minute incubation in between. Eluted proteins and the flow-through were combined and concentrated by 10 kDa Amicon Ultra-15 centrifugal filters (Millipore, #UFC901008). The sample was buffer exchanged once into the wash buffer (+ 20 μM Shield-1) using the same concentrator to remove 90% of the DTT in the solution. Protein concentration was determined by absorption at 488 nm using the extinction coefficient of sfGFP[5] as 83300 $M^{-1}cm^{-1}$. Approximately 5 mg of AgDD was obtained from 1 L of culture. Proteins were aliquoted and stored at 200-400 μM at -80 °C. Multiple freeze-thaw cycles were avoided.

Ubiquitin chains were synthesized using human ubiquitin mutant $Ub^{L73F}$ to minimize nonspecific deubiquitylation in the extract[6]. $Ub^{L73F}$ with a cysteine residue and a 6xHis-tag at the N-terminus was purified from *E. coli* and labeled with Dylight-550-maleimide (Pierce, #62290) as described previously[7]. Unreacted cysteines of $Ub^{L73F}$ were alkylated by N-Ethylmaleimide. Human E1 and E2-25K were purified from *E. coli*[7]. Uev1a and UbcH13 were purchased from R & D Systems.

### HeLa S3 Cell Extract Preparation

HeLa S3 extract was prepared as described previously[8]. Briefly, 2 L of spinner culture of asynchronous HeLa S3 cells were collected and homogenized using nitrogen cavitation in the swelling buffer (20 mM HEPES pH 7.5, 5 mM KCl, 1.5 mM $MgCl_2$, 1 mM DTT, 1× protease inhibitor cocktail). The homogenate was sequentially centrifuged at 5000×g for 10 minutes and 20000×g for 30 minutes. The supernatant was collected, aliquoted, and stored at -80 °C.

### AgDD Aggregates Formation

Purified AgDD stabilized by Shield-1 was centrifuged at 17000×g for 15 minutes at 4 °C before use. The supernatant was added to *Xenopus* egg extract or a physiological buffer (1 mM DTT, 100 mM $KCH_3COOH$, 30 mM KCl, 1 mM $MgCl_2$, 1 mM $Na_2ATP$, 10 mM $Na_2HPO_4$) at a 10 μM final concentration in the presence of 200 μM FKBP(F36V) (or 20 μM Shield-1 for uninduced controls) and incubated at 18 °C. Samples were taken every 10 minutes after 30-minute incubation and examined on microscope slides using an epi-fluorescent microscope equipped with a GFP channel filter and a 10x objective. Aggregates were allowed to form for an additional 30 minutes after they were first visible under the scope. Samples were then kept on ice and used for experiments on the same day.

### Synthesis of Fluorescent Ubiquitin Chains

Ubiquitin chains with K48 or K63 linkage were synthesized in reactions containing 100 μM $Ub^{L73F}$ and 1 μM DyLight 550-labeled $Ub^{L73F}$, 0.2 μM E1, 15 μM E2-25K (for K48 linkage) or 5 μM Uev1a/UbcH13 (for K63 linkage) in the buffer (15 mM HEPES pH 7.5, 10 mM ATP, 10 mM $MgCl_2$, 0.6 mM DTT, and 1 mM PMSF); and incubated at 37 °C for 5 hours. The product was buffer-exchanged using a Zeba 40K column (Thermo Scientific, #87764) into PBS. Protein concentration was determined by SDS-PAGE and Coomassie staining using BSA as a concentration standard.

### Conjugation of Ubiquitin Chains to Dynabeads

Dynabeads M-270 beads with amine groups were washed twice and resuspended by the freshly prepared passivation buffer (500 mM $K_2SO_4$, 100 mM $NaHCO_3$). We used 5 kDa PEG plus 2.5% 5 kDa biotin-PEG (LaysanBio, mPEG-SVA-5000; Biotin-PEG-SVA-5000) in a "clouding-point" solution as described previously[8] to passivate the beads for 2 hours at 25 °C with slow tilt and rotation. Beads were then washed twice with TBS with 0.5 mg/mL BSA, 0.01% Tween-20, and resuspended to the original volume. 5 μL of beads were then incubated with 8 μg of Streptavidin (Invitrogen, #434301) for 30 minutes at 25 °C and washed three times with TBS.

To conjugate the ubiquitin chains to the beads, 10 μL solutions of 50 μM ubiquitin chains or ubiquitin monomers were biotinylated by 100 μM NHS-PEG4-biotin (Thermo Scientific, #21363) at 25 °C for 1 hour and buffer exchanged twice into PBS using a Zeba 7K column. Samples were then incubated with 5 μL streptavidin-coated Dynabeads M-270 beads at 25 °C for 1 hour and washed three times with PBS. 0.5 μL of beads were boiled and blotted by Alexa Fluor 790-streptavidin (ThermoFisher, #S11378). Biotin-XX goat anti-mouse IgG secondary antibody (ThermoFisher, #B2763) with a known concentration (2 mg/mL) and degree of biotin-XX labeling (4 biotins per antibody) was used as the standard. Surface density of ubiquitin on the beads was estimated as below:

$$\text{\# of ubiquitins per bead} = \frac{\text{intensity of Ub bands}}{\text{intensity of Biotin}-\text{XX antibody band}} \times \frac{4 \times \text{\# of molecules of the antibody}}{2e9 \text{ beads }/\text{mL} \times 0.5\ \mu\text{L}};$$

$$\text{surface coverage rate of ubiquitins on the bead} = \frac{\text{\# of ubiquitins per bead} \times \pi \times 1.5\ \text{nm}^2}{4\pi \times 1.4\ \mu\text{m}^2}.$$

The labeling protocol yielded on average one biotin molecule per ubiquitin. Given the bead diameter as 2.8 μm and ubiquitin molecule diameter as 3 nm, the surface coverage rate of the beads by ubiquitin chains or ubiquitin monomers was estimated to be 5%.

### Cell Line Construction and General Culture Conditions

WT HEK293T and U2OS cells were obtained from ATCC and cultured in DMEM supplemented with 10% heat-inactivated FBS and 2× Antibiotic-Antimycotic (ThermoFisher, #15240062) at 37 °C and 5% $CO_2$. AgDD-sfGFP, AgDD-sfGFP-HOOK2, and AgDD-sfGFP-HOOK3 and Htt (Q94)-CFP were integrated into HEK293T or U2OS cells by lentivirus, followed by blasticidin selection (InvivoGen, #ant-bl-05) for two weeks to make stable cell lines. Cells transduced with the lentivirus carrying H2B-mCherry were sorted on a Beckman Coulter MoFlo Astrios EQ high-speed cell sorter to obtain pure populations expressing mCherry. Lentivirus was prepared following a standard protocol: viruses were collected from 10 cm dish 48 hours post transfection of the transfer vector plasmid, packaging plasmid (Addgene plasmid #8455) and envelope plasmid (Addgene plasmid #8454) at a ratio of 4:2:1 and were concentrated to 600 μL by 100 kDa Amicon Ultra-15 centrifugal filters (Millipore, #UFC910008) after centrifugation at 1500×g for 20 minutes at 4 ℃. 200 μL of virus solution was added to 1 well of HEK293T or U2OS cells seeded in a 12-well plate at 60% confluency. Cells were moved to larger containers after 24 hours. Cells were maintained in the same media with 1 μM Shield-1 (AOBIUS, #AOB1848). U2OS cells co-expressing AgDD-sfGFP and HaloTag-H2A were generated together using a lentiviral system (AgDD-sfGFP) and a retroviral system (HaloTag-H2A)[1] and selected using blasticidin and hygromycin (ThermoFisher, #10687010). Transient transfection of the Htt (Q94)-CFP construct into HEK293T cells was performed using TransIT-293 Transfection Reagent (Mirus Bio, #MIR 2700) according to the manufacturer's instructions. 1 μg/mL doxycycline (Takara Bio, #631311) was used for inducing expression from the TetOn promoter.

AgDD aggregation was induced by adding purified recombinant FKBP(F36V) (1.75 mM stock solution in PBS plus 10% glycerol) to the culture at a final concentration of 5 μM to deplete free Shield-1. Effects of drugs were tested by pretreating the cells with each drug for 30 minutes before aggregation induction, and drugs were kept in the media during the experiment.

siRNAs targeting human CCT2, CCT4, CCT8, HDAC6, BAG3, and the negative control (NC) were purchased from IDT (DsiRNAs) and transfected to HEK293T cells using Lipofectamine™ RNAiMAX Transfection Reagent (Invitrogen, #13778150) at a final concentration of 20 nM. For double-KD of two genes, each siRNA was used at 10 nM. Cells were incubated at 37 ℃ for 24/48/72 hours before being harvested for immunoblotting. Two siRNAs were tested for each target gene, and one was selected for functional tests, where cells were induced to form the aggresome 48 hours after transfection of the siRNAs, and the time required to form a single punctum of the aggresome was measured through live-cell imaging.

### Quantification of the Intracellular AgDD Concentration

A calibration curve between sfGFP fluorescent intensity and concentration was obtained for purified sfGFP protein using a TECAN Infinite F200 fluorescence microplate reader with a GFP filter (EX 485/20; EM 516/20). AgDD-sfGFP signal in the cell suspension (in PBS) was measured and converted to the molar concentration ($[\mathrm{AgDD}]_{\mathrm{PBS\ suspension}}$) after multiplying by the coefficient determined as the slope of the calibration curve. sfGFP signal was mostly from full-length AgDD-sfGFP, as shown by immunoblotting (Fig. S1B). The average intracellular AgDD concentration was calculated as:

$$[\mathrm{AgDD}]_{\mathrm{intracellular}} = \frac{[\mathrm{AgDD}]_{\mathrm{PBS\ suspension}} \times \mathrm{Volume\ of\ PBS\ suspension}}{\mathrm{Number\ of\ 293T\ in\ PBS\ suspension} \times \frac{4}{3}\pi R^3} \quad (1)$$

where $R$ is the radius of the HEK293T cell, set at 5 μm.

To estimate the molar concentration of AgDD in each single cell from microscopy data, 100 unperturbed cells were randomly selected from 5 fields-of-view and manually segmented to determine the cytoplasmic sfGFP signal ($\mathrm{sfGFP}_{\mathrm{single\ cell}}$). The average cytoplasmic sfGFP signal ($\mathrm{sfGFP}_{\mathrm{bulk}}$) was calculated as the mean of $\mathrm{sfGFP}_{\mathrm{single\ cell}}$. The molar concentration of AgDD in each cell was calculated using the value from (1):

$$[\mathrm{AgDD}]_{\mathrm{single\ cell}} = \frac{\mathrm{sfGFP}_{\mathrm{single\ cell}} \times [\mathrm{AgDD}]_{\mathrm{intracellular}}}{\mathrm{sfGFP}_{\mathrm{bulk}}}. \quad (2)$$

### Immunofluorescence Staining

HEK293T or U2OS cells expressing AgDD-sfGFP were seeded on Poly-L-Lysine-coated (Millipore Sigma, #P4707-50ML) coverslips 24 hours before experiments. 4 hours after aggresome induction, cells were washed once with cold phosphate-buffered saline (PBS), fixed in 4% paraformaldehyde (Leagene Biotechnology, # DF0135) for 15 minutes at 25 °C, washed 3×5 minutes in PBS, permeabilized with PBS + 0.2% Triton X-100 for 10 minutes, and blocked for 30 minutes in PBS + 3% BSA, 0.1% Triton X-100. Centrosomes were stained by incubating overnight with 1:500 diluted rabbit polyclonal pericentrin antibody (Abcam, #AB4448) in blocking buffer at 4 °C. Vimentin was stained by incubating overnight with 1:200 diluted mouse monoclonal vimentin antibody (Abcam, #8069). Cells were then washed 3×5 minutes in PBS + 0.1% Triton X-100, incubated with 1:1000 dilution secondary antibodies (Donkey-anti-Rabbit 647 (Invitrogen, #A32795); Donkey-anti-Mouse 568 (Invitrogen, #A-10037)) in blocking buffer for 1 hour at 25 °C, washed 3×5 minutes in PBS, and mounted on microscope slides using ProLong Gold Antifade Mountant with 4′,6-diamidino-2-phenylindole (Invitrogen, #P36930) and imaged the following day. Images were taken using the same Nikon spinning-disk confocal microscope for live-cell imaging with a Nikon CFI Apo TIRF 60x Oil objective.

To study colocalization of aggresome and the chaperonin complex, HEK293T cells transfected with the Htt (Q94)-CFP construct were induced with 1 μg/mL doxycycline for 12 hours before fixation. Cells were co-stained with 1:200 diluted mouse monoclonal antibody for mono- and poly ubiquitinylated proteins (FK2, Sigma-Aldrich, #04-263), 1:50 diluted rabbit polyclonal CCT2 antibody (Proteintech, #24896-1-AP), and 1:100 diluted rabbit polyclonal CCT8 antibody (Proteintech, #12263-1-AP) for 2 hours at 25 °C.

### Live-Cell Imaging

Cells were seeded onto a glass-bottom 8-well µ-Slide (ibidi, #80821) or glass-bottom 12-well plate (Cellvis, #P12-1.5H-N) coated with fibronectin (Sigma-Aldrich, #F0895) one day before the experiment in regular DMEM medium without phenol red (Cytiva, #SH30284.01) plus 10% heat-inactivated FBS and 2× Antibiotic-Antimycotic. Cells expressing HaloTag-H2A were stained with 500 nM JFX650-HaloTag ligand (Janelia) for 2 hours right before imaging. Widefield fluorescence microscopy was performed on a Nikon Ti2 inverted microscope with epi-fluorescence optics, Lumencor Spectra-X light engine for fluorescence illumination, a sCMOS camera (Hamamatsu Flash4.0, 6.5 $\mu m^2$ photodiode) and a microscope incubator (OkoLab) for controlling temperature (37 °C), humidity, and $CO_2$ level (5%), and 20x (Nikon Plan Apo 0.75 NA) and 40x (Nikon Plan Apo 0.95 NA) objectives. Images were taken every 10 minutes. NIS Elements software was used for image acquisition. Cells were covered with mineral oil (Sigma, #M8410) to prevent evaporation. Confocal microscopy was performed on a Nikon spinning-disk confocal microscope (Nikon Ti2 inverted microscope, Yokogawa CSU-W1) with 405/488/561/640 nm lasers (iChrome MLE), a sCMOS camera (Photometrics Prime 95B), a Nikon CFI Plan Apochromat VC 60x water immersion objective, and a microscope incubator (OkoLab). Images were taken every 1 minute for single particle tracking. Images were acquired using the NIS Elements software and examined using ImageJ (version 1.53t, National Institutes of Health).

To test the aggresome formation efficiency in HEK293T cells expressing Htt (Q94)-CFP with or without chaperonin knockdown, cells were seeded on a glass-bottom 12-well plate and imaged using the IncuCyte® ZOOM Live-Cell Analysis System (Sartorius) with a 20x objective and phase and GFP filters. Images were taken every 30 minutes for 60 hours.

### Cargo Transport Assay in *Xenopus* Egg Extract (XE)

#### XE Preparation

Actin-fragmented mitotic *Xenopus* egg extract (CSF extract) was used in all egg extract experiments except the one shown in Supplementary Video 4. Actin-fragmented CSF extract was either prepared with intact actin as described[9] and treated with 20 µg/mL cytochalasin D (CytoD) right before experiments or prepared from 100 µg/mL CytoD-treated *Xenopus* eggs and supplemented with 10 µg/mL CytoD[10] while otherwise following the same protocol[9]. Freshly prepared CSF extract was kept on ice and used for experiments within 8 hours. Freshly prepared CSF extracts could also be aliquoted, slowly cooled in a Mr. Frosty™ Freezing Container (Thermo Scientific, #5100-0001), and stored at -80 °C. Frozen aliquots of egg extract were thawed on ice and used for later experiments while avoiding multiple freeze-thaw cycles. Whenever possible, extracts were pipetted with 200 µL wide bore tips (Axygen, #T205WBC) to reduce shear damage. We compared behaviors of aggregate transport using freshly prepared XE, frozen XE, XE made with CytoD, or XE made with intact actin but treated with CytoD right before the experiment and observed consistent trends of positive size selectivity (Fig. S7).

Actin-intact cycling egg extract used in Supplementary Video 4 was prepared largely following Guan, et al., 2018[11] without adding CytoD. Instead of enclosed imaging chambers (described below), cycling extract was loaded onto a passivated coverslip and covered with mineral oil for imaging.

#### Cargo Transport Assay

To assemble and image interphase microtubule asters in egg extract, actin-fragmented CSF extract was supplemented with fluorescent tubulin to label microtubules, cycled to interphase to allow aster growth, and supplemented with demembranated *Xenopus* sperms to nucleate asters. Tubulin purified from bovine brain was labeled with Alexa Fluor 647 (tubulin-Alexa647) as previously described[12]. In a typical reaction, tubulin-Alexa647 was added to CSF extract on ice to a final concentration of 50 nM. To trigger exit from CSF arrest and entry to interphase, $CaCl_2$ was then added to a 0.4 mM final concentration. The extract was mixed well immediately after calcium addition by gently flicking and incubated in an 18 °C water bath for 4 minutes, then returned to ice for 3 minutes. Demembranated *Xenopus* sperms were prepared as described[13] and added to the appropriate concentration that minimized contact between asters. For the experiment shown in Supplementary Video 4, EB1-mApple[14] was added to a final concentration of 200 nM to label the centrosomes.

To study dynein-dependent transport, AgDD aggregates (see above), dynein-adaptor-coated beads (see below) and unpassivated carboxylate beads (Polysciences, #21636; Fig. S6) were 1:20 diluted into the working extract and mixed well by flicking the tube. The extract reaction was loaded into customized imaging chambers assembled from coverslips passivated with poly-L-lysine-g-polyethylene glycol (PLL-g-PEG), as described previously[9, 15]. Imaging was started immediately using a Nikon Eclipse Ti2-E inverted microscope with SOLA SE V-nIR light engine and an Andor Zyla 4.2 PLUS sCMOS camera. The microscope room was kept at 18 °C. Low-temporal-resolution tracking experiments were performed using a Nikon CFI Plan Apo Lambda 10x NA 0.45 objective lens and imaged every one minute unless otherwise stated. High-temporal-resolution tracking was performed using a Nikon CFI Plan Apo Lambda 40x NA 0.95 objective lens and imaged at 30 Hz. Two fluorescence channels were used in imaging, including the GFP channel for dynein cargoes (Filter cube: Nikon 96362/CHROMA 49002 ET GFP, excitation filter 470/40, emission filter 525/50) and the Cy5 channel for microtubules (Filter cube: CHROMA 49009 ET Cy5 NX, ex 640/30, em 690/50). In experiments with multiple conditions imaged in parallel, the slide holder was first chilled on ice for several seconds, so that aster growth would start at the same time across all conditions.

To inhibit dynein, the p150-CC1 fragment of dynactin[16] was added to a final concentration of 40 μg/mL.

To test the deubiquitylation activities of Usp2 in the extract, 1 μM Usp2 catalytic domain $Usp2^{CD}$ (R & D Systems, stock solution at 50 μM in 50 mM HEPES pH 8.0, 150 mM NaCl, 0.1 mM EDTA, 1 mM DTT) or an equal volume of the same buffer was added to the $Ca^{2+}$-activated extract and incubated at 18 °C. Samples were taken every 10 minutes during the incubation and immunoblotted with an antibody for ubiquitin (Cell Signaling, #43124). To remove any ubiquitin on the AgDD aggregates formed in the extract, the aggregates were incubated with 10 μM $Usp2^{CD}$ at 18 °C for 10 minutes before being diluted into the extract for the transport assay.

**Conjugation of Dynein Adaptors to Dynabeads**

Dynabeads Protein G beads (Thermo Fisher, #10004D) were washed twice and resuspended in PBS to 10-fold bead volume (BV), then incubated with 5-10 μg of primary antibody per 20 μL beads (initial BV) overnight at 4 °C with slow tilt and rotation. Beads were washed with 10× BV of PBS to remove unbound antibodies and resuspended to 1/5 of the initial BV. Antibodies targeting human proteins HDAC6 (Proteintech, #12834-1-AP), CCT8 (Proteintech, #12263-1-AP), 26S proteasome (MCP21; Enzo Life Sciences, # BML-PW8105), Hsc70 (Proteintech, #10654-1-AP), Hsp90 (Proteintech, #13171-1-AP) and p62 (BD Biosciences, #610832), as well as random rabbit IgG (Jackson ImmunoResearch, #011-000-003) were purchased. The antibody targeting *Xenopus* HOOK2 was made and purified from rabbit serum by the Mitchison lab[15]. To immunoprecipitate human adaptor proteins (except HOOK2) from HeLa S3 extract, 20 µL beads coated with primary antibodies were incubated with 100 µL HeLa cell extract diluted with 200 µL of TBST (TBS + 0.05% Tween-20) with 1× protease inhibitor cocktail for 2 hours at 4 ℃ and washed three times with TBST. Beads were then washed twice, resuspended, and stored in a physiological buffer (1 mM DTT, 100 mM $KCH_3COOH$, 30 mM KCl, 1 mM $MgCl_2$, 1 mM $Na_2ATP$, 10 mM $Na_2HPO_4$). For preparing HOOK2-coated beads, 20 µL beads were incubated with 100 µL CSF extract (actin depolymerized, $Ca^{2+}$ activated) for 60 minutes at 4 °C, washed 5 times with the wash buffer (20 mM KCl, 1 mM $MgCl_2$, 10 mM K·HEPES pH 7.7, 1 mM EGTA) and resuspended and stored in the physiological buffer. Right before the transport assay, fluorescent secondary antibody was added to dynein adaptor-coated beads to a final concentration of 20 µg/mL, including Goat anti-Rabbit IgG Secondary Antibody Alexa Fluor™ 488 (Invitrogen, #A-11008) for rabbit-derived primary antibodies and Goat anti-Mouse IgG Secondary Antibody Alexa Fluor™ 488 (Invitrogen, #A-11001) for mouse-derived primary antibodies. Beads were validated by mass spectrometry as described below.

**Immunoprecipitation of AgDD Interacting Proteins**

Dynabeads Protein G beads were saturated with anti-HA primary antibodies (HA-7; Sigma, #H9658) as described previously. An extra step of crosslinking primary antibodies onto the beads was performed to minimize interference detection of antibodies in the mass spectrometry analysis. Beads were washed twice with 20× bead volume (BV) of 0.2 M sodium tetraborate pH 9.0 and resuspended in 10× BV of the same buffer. Dimethylpimelimidate was weighed and added as powder to reach a final concentration of 20 mM and dissolved by gentle mixing. Reactions were rotated for 30 minutes at 25 ℃ and quenched by washing the beads twice with 10× BV of the quenching buffer (150 mM NaCl, 200 mM Tris-HCl pH 7.5), with a 10-minute incubation in between. Beads were then washed twice with the wash buffer (0.1 M KCl, 10 mM HEPES pH 7.7, 100 µg/mL BSA) and resuspended to 1/5 of the original volume. 4 µL of concentrated bead slurry was used to immunoprecipitate proteins from 100 µL of extract.

Two batches of actin-depolymerized CSF extracts were freshly prepared in parallel and used for experiments as repeats. 15 µL AgDD aggregates formed in the extract ($AgDD_{XE}$), in the physiological buffer ($AgDD_{pb}$) and the uninduced control ($AgDD_{sol}$) were prepared as described previously. Both extracts were activated by $Ca^{2+}$ and each was split into 4×2 mL Protein LoBind Tubes (Eppendorf, #022431102), 100 µL each. 10 µL $AgDD_{XE}$, $AgDD_{pb}$, $AgDD_{sol}$, or 10 µL extract was added to each tube. 4 µL HA-antibody-coated beads were then added to each tube and mixed well by flicking. Samples were incubated for 45 minutes at 16 ℃ with slow tilt and rotation. Beads were then collected on a magnet for 5 minutes on ice and washed five times with 1 mL wash buffer (25 mM KCl, 10 mM K·HEPES pH 7.7, 1 mM $MgCl_2$, 1 mM EGTA, 1 mM ATP). Beads were transferred to a new tube twice after the 4th and 5th washes.

Immunoprecipitated proteins were identified by mass spectrometry as described below.

### Quantitative Mass Spectrometry

#### Sample Preparation

Proteins immunoprecipitated by the beads were determined by Tandem-Mass-Tag mass spectrometry (TMT-MS). On-bead digestion was performed by resuspending beads in 1× BV of

50 mM EPPS pH 8.3 and 2 M GuHCl with 1:100 diluted (v/v) lys-C (Promega, #VA1170; 2 mg/mL stock solution). The reactions were incubated at 37 ℃ for 4 hours on an Eppendorf ThermoMixer C at the top speed. One same dose of lys-C was added again, and the reactions were incubated overnight. Supernatant was transferred to a new tube and diluted to 4× BV with 50 mM EPPS pH 8.3. 1:50 diluted (v/v) trypsin (Promega, #V511C; 0.5 mg/mL stock solution) was added to the sample and incubated at 37 ℃ for 4 hours on a tube rotator. The peptide concentration was determined using CBQCA fluorescent assay following the manufacturer's instruction. TMT labeling, alkylation/cysteine protection by iodoacetamide (IAA), stage tip desalting, and SpeedVac drying were performed following the manufacturer's instructions and standard protocols (ThermoFisher, #A44520; Zhou, et al., 2024[2]). Peptides were reconstituted in 0.1% formic acid for LC-MS analysis. TMT-labeling efficiency was verified to be more than 95%.

**LC-MS Data Collection and Analysis**

*Samples from Dynein-Adaptor-Coated Beads*

Mass spectrometric data were collected on an Orbitrap Exploris480 instrument. The mass spectrometer was coupled to a Proxeon NanoLC-1200 UHPLC attached to a 100 µm capillary column packed with 35 cm of Accucore 150 resin (2.6 µm, 150Å; ThermoFisher Scientific) at a flow rate of ~420 nL/min. Data were acquired using multiple injections (n=2) with varying combinations of FAIMS compensation voltages (CVs) between -30 and -80 V (3 CVs per set) over a 150 min gradient. A 1-second TopSpeed cycle was used for each CV. The scan sequence began with an MS1 spectrum (Orbitrap analysis, resolution 60000, 350-1350 Th, automatic gain control (AGC) target 100%, maximum injection time "auto"). The hrMS2 stage consisted of fragmentation by higher energy collisional dissociation (HCD, normalized collision energy 32%) and analysis using the Orbitrap (AGC "standard", maximum injection time 96 ms, isolation window 0.7 Th, resolution 45000).

The acquired data were searched using the open-source Comet algorithm (release_2019010)[17-21]. Spectral searches utilized a custom FASTA-formatted database containing common contaminants and reversed sequences (Uniprot Human, 2021). The following parameters were used: 50 PPM precursor tolerance, fully tryptic peptides, a fragment ion tolerance of 0.02 Da, and a static modification by TMTPro16 (+304.2071 Da) on lysine and peptide N-termini. Additionally, carbamidomethylation of cysteine residues (+57.0214 Da) was applied as a static modification and oxidation of methionine residues (+15.9949 Da) as a variable modification. Peptide spectral matches (PSMs) were filtered to achieve a peptide false discovery rate (FDR) of 1% using linear discriminant analysis and a target-decoy approach. Resulting peptides were further refined to reach a final protein-level FDR of 1% at the dataset level, with proteins grouped accordingly. Reporter ion intensities were adjusted for synthesis impurities of different TMT reagents as specified by the manufacturer. For each MS2 spectrum quantification, a minimum total signal-to-noise (S/N) of 100 or 160 for all reporter ions was required depending on the experiment. Finally, protein abundance measurements were normalized so that the total signal-to-noise across all channels for a given protein was set to 100, thus providing a measure of relative abundance.

*Samples from AgDD-Coated Beads*

Mass spectrometric data were collected on an Orbitrap Fusion Lumos instrument (using hrMS2-mode). The mass spectrometer was coupled to a Proxeon NanoLC-1200 UHPLC attached to a

100 µm capillary column packed with 35 cm of Accucore 150 resin (2.6 µm, 150Å; ThermoFisher Scientific) at a flow rate of ~440 nL/min. Data were acquired using multiple injections (n=3) with varying combinations of FAIMS compensation voltages (CVs) between -30 and -80 V (3 CVs per set) over a 150 min gradient. A 1-second TopSpeed cycle was used for each CV. The scan sequence began with an MS1 spectrum (Orbitrap analysis, resolution 60000, 350-1350 Th, automatic gain control (AGC) target 100%, maximum injection time "auto"). The hrMS2 stage consisted of fragmentation by higher energy collisional dissociation (HCD, normalized collision energy 36%) and analysis using the Orbitrap (AGC 300%, maximum injection time 250 ms, isolation window 0.7 Th, resolution 50000). The acquired data were analyzed as before.

**Identification of AgDD Interactome**

Among all conditions ($AgDD_{XE}$, $AgDD_{sol}$, $AgDD_{pb}$, naked beads), the enrichment of proteins captured by $AgDD_{XE}$ compared to $AgDD_{sol}$ was used to identify candidate proteins potentially involved in aggresome formation. For each identified protein, the mean of the $AgDD_{XE}$ to $AgDD_{sol}$ ratio of two repeats was calculated. Proteins were selected if the mean ratio was greater than 1.15. Gene Ontology (GO) analysis was performed on the selected genes using shinyGo[22] based on the category of Molecular Function[23, 24].

### Microscopy Image Analysis

#### Live-Cell Image Analysis

*Aggresome Detection*

AgDD aggresome formation is operationally defined as the appearance of a perinuclear punctum if the surrounding aggregates move toward it in subsequent frames and all peripheral aggregates are eventually sequestrated into it. The aggresome-containing cells were identified manually. Htt (Q94) aggresomes were identified using the ImageJ plugin "*TrackMate*."[25] Briefly, a stable HEK293T cell line integrated with pTetOn-Htt (Q94)-CFP was transfected with CCT2-targeting siRNA and imaged using the IncuCyte® ZOOM System. Doxycycline was added to the media to induce Htt (Q94) expression at different time points as indicated in Fig. S10E-F. Cell confluency was monitored via the IncuCyte® ZOOM Confluence Processing analysis tool (Basic Analyzer) based on the phase channel. The number of aggresomes in each frame was determined by *TrackMate.* Since most cells contained a single Htt (Q94) aggresome, the fraction of aggresome-containing cells at each time point was calculated as the ratio of the aggresome number to the total cell number. The latter was inferred from the confluency value based on a calibration using manually-counted cell number vs. confluency in 10 fields.

*AgDD Aggresome Enrichment Factor*

Aggresome enrichment factor was defined as the maximal fluorescence intensity of pixels in the perinuclear region ($\boldsymbol{I}_{max}$) divided by the average pixel intensity of the whole cell ($\boldsymbol{I}_0$), before Shield-1 removal. The ring-shaped perinuclear region with a width of ~2 µm was manually segmented based on the contour of the nucleus. For aggresome-containing cells, the width was enlarged to enclose the entire aggresome.

#### Single-Particle Tracking of AgDD Aggregates

*MTOC Localization*

For low-resolution tracking experiments in *Xenopus* egg extract (XE), the MTOC was manually registered as the center of the aster based on Alexa Fluor 647 signal using the ImageJ plugin "*Manual Tracking*." For high-resolution tracking experiments in XE and on live cells, the MTOC was determined as the center of the aggregates that were already clustered around the centrosomes through the *TrackMate*. Clustered aggregates were first identified via "LoG detector" as a single object, whose diameter was adjusted to tightly enclose all the clustered aggregates. The object was then tracked via "LAP Tracker" with appropriate frame-to-frame linking and gap closing based on visual examination. The center of the object was used as the MTOC position. An extra step of aligning all frames through centering the MTOC was performed before tracking the aggregates for the low-resolution experiments in XE, to correct for the common drift of asters during the time-lapse imaging lasting for tens of minutes.

*Aggregates Tracking*

Aggregates were tracked using *TrackMate*. Aggregates were first identified via "LoG detector" with appropriate parameters:

- "Estimated objective diameter": adjusted to the minimal value that could cover all visible aggregates, named $r_0$ which will be used for size determination below.
- "Quality threshold": adjusted based on visual examination.
- "Pre-process with median filter" and "Sub-pixel localization" were both selected.

Identified aggregates were then tracked via "LAP Tracker" with appropriate parameters:

- "Frame-to-frame linking": "Max distance" was adjusted to be slightly larger than the largest displacement detected between consecutive frames; "Feature penalties" included "Mean intensity ch1 (weight=1)" and "Quality (weight=1)."
- "Track segment gap closing": "Max frame gap" was set to "1" and "Max distance" was chosen to be the same as "Max distance" in "Frame-to-frame linking."

Lastly, after removing mis-tracked trajectories manually, the $(x, y)$ positions of each trajectory at each time point $t$ were saved and converted to the Euclidean distance $d(t)$ to the MTOC in MATLAB (version R2023a, The MathWorks, Natick, MA). The "trajectory" over time was calculated as $d(t=0) - d(t)$, the "travel distance towards MTOC," unless otherwise stated. A positive value means moving towards the MTOC.

*Size Determination via 2D Gaussian Fitting*

For each timepoint from each trajectory, a $(2r_0+1) \times (2r_0+1)$ square centered at the $(x, y)$ position of the aggregate was extracted from the original image and fitted by a 2D Gaussian function[26] in MATLAB using the following equation:

$$I(a,b) = p_0 + p_1 \exp\left[-\left(\frac{(a-p_5)\cos p_2 + (b-p_6)\sin p_2}{p_3}\right)^2 - \left(\frac{-(a-p_5)\sin p_2 + (b-p_6)\cos p_2}{p_4}\right)^2\right],$$

where $a$ and $b$ are the coordinates in pixel, $I(a,b)$ is the intensity of the pixel, $p_{0-6}$ are the parameters to fit. To be noted, $p_3$ and $p_4$ are restricted to positive values and $p_2$ is restricted to be between 0 and 180 degrees without loss of generality. Other parameters constraining the fitting algorithm were defined in the script. Aggregates with irregular shapes or a low signal-to-noise ratio were discarded, if satisfying any of the following conditions:

i. $p_5$ or $p_6$ is smaller than 1 or larger than $(2r_0+1)$.
ii. $p_3$ or $p_4$ is negative.
iii. $p_1$ is smaller than $p_0$.

The diameter of the aggregate was calculated as $c_0\sqrt{p_3 p_4}$, i.e., $2c_0\sqrt{\sigma_X \sigma_Y}$ where $\sigma_X$ and $\sigma_Y$ are the standard deviations (i.e., spread parameters) along the *x* and *y*-axis. $c_0$ was determined from a calibration curve using standard carboxylate beads of known sizes (Polysciences, #21636; Fig. S3).

*Determination of the Localization Error in the High-Resolution Tracking Experiments*
The same bead standards used for size calibration were immobilized on the coverslip via nonspecific attachment and imaged using the same settings as in the high-resolution tracking experiments for AgDD aggregates and adaptor-coated beads (Fig. 3, 4C-E). Data were then analyzed by *TrackMate* as described before. The localization error was calculated as the root-mean-square deviation (RMSD) of the bead's position after subtracting the drift, which was determined as the linear regression line of the position over time.

**Trajectory Segmentation and Diffusion Constant Calculation**
Segmentation was performed using a MATLAB package "*slmengine*"[27] with a fixed root mean squared error (RMSE) or a fixed number of segments. For low-resolution tracking experiments, we used a fixed RMSE = 0.025 and set the maximal segment number as 10. For high-resolution experiments, we first used a fixed RMSE = 0.01 and set the maximal segment number as 20 to process the entire trajectory, and then extracted the transport segments determined as described below to perform a second round of segmentation using a fixed segment number, calculated as the transport segment length in seconds divided by 0.5. All parameters mentioned above were chosen based on visual examination.

For each trajectory, the velocity of each segment was determined as the slope after performing linear regression on the distance over time. Segments with the smallest absolute values of velocity were selected, which were most likely to be the "Pause" state. The number of segments selected as pauses was less than 1/3 of the total segment number. Selected pauses were then used to calculate the mean square displacement (MSD) over time and fitted by a linear function. The diffusion constant was calculated as half of the slope. The mean value of the diffusion constant calculated from all selected segments was used as the diffusion constant *D* for the trajectory. A hypothesis test for all other segments not selected as pauses was then performed, given the null hypothesis that "*the displacement given the segment length t should follow a Gaussian distribution with the mean equal to 0, and the standard deviation equal to the square root of 2Dt.*" The segment was designated as "Transport engaged (+)" for low-resolution experiments and "Transport (+)" for high-resolution experiments if the displacement fell to the right tail of the Gaussian distribution; and "Transport engaged (-)" or "Transport (-)" for the left tail. For the low-resolution experiments, the cutoff bounds on both tails were 0.025, achieving a goodness-of-fit = 0.072 (normalized root mean square error (NRMSE), calculated as RMSE divided by the range of the variable). For high-resolution experiments, the cutoff bounds are 0.05 on both tails for the first round of segmentation and 0.005 and 0.1 on the left and right tails respectively for the second round of segmentation, achieving a goodness-of-fit = 0.0217 (NRMSE). These parameters were chosen to allow the segmentation results most consistent with visual examination. Lastly, any adjacent segments with the same designations were connected as a single segment.

**Determination of the Aggregate's Velocities**
*MTOC-Directed Frame-by-Frame Velocity* (Fig. 1J, 5D, S7, S8):

For each aggregate particle, the distance between the MTOC and the aggregate at each frame was calculated. The MTOC-directed velocity of the aggregate was determined as the change in the distance between two consecutive frames divided by the time interval in between, with a positive value meaning movement towards the MTOC. For each particle, the velocity at each time point was associated with the particle size determined at the same time point.

*Time-Averaged Transport Velocity of Single Particle* (Fig. 2D, 3A, S6):
When the aggregate size remains largely stable during the time-lapse imaging (Fig. S5), the transport velocity and particle size were determined as the time average over the entire trajectory. Specifically, the transport velocity of each aggregate particle was calculated as the total distance moving towards the MTOC divided by the duration of the trajectory, with a positive value meaning movement towards the MTOC.

*Instantaneous Velocity* (Fig. 3C, 3D):
The instantaneous velocity was determined as the time derivative of the trajectory. A 2-second or 0.5-second time window was used for high-resolution tracking data as indicated.

*Intrinsic Velocity* (Fig. 3E, 3F, 4E):
For each particle, the intrinsic velocity was defined as the mean of the top 20% values of the instantaneous velocity within transport segments, unless otherwise stated.

**Violin Plot**
All violin plots were generated using a MATLAB function "*violin*"[28] with additional modifications.

### A Physical Model for Dynein-Mediated Transport of Protein Aggregates

Our goal here is to establish a physical model of dynein-mediated cargo transport and determine how the average transport velocity depends on the system parameters. The entities in the model are dynein (Dy), microtubule (MT), protein aggregates or cargo (C), and their complexes. The physical processes being modeled are binding-unbinding between the model entities and the viscosity-limited transport by dynein. The overall procedure is to derive the system partition function which dictates the likelihoods of C in different states. The likelihood of C in the active transport state directly determines the average transport velocity. Although dynein transport is a non-equilibrium process, we assume it is transient, allowing us to treat binding and unbinding events as if they were in equilibrium.

The cargo was identified with two states in our experiments: a freely diffusing state and a local state (denoted by the subscript "L") that is close to MT. Cargoes starting from a freely diffusing state, either staying alone (C) or bound with free dynein (C-Dy), must first approach MT, i.e., enter the local state, before forming an active transport complex ($C_L$-Dy-MT) and engaging in transport. We do not pre-define a transition sequence and the cargo can jump freely among possible states as depicted in the figure on the right.

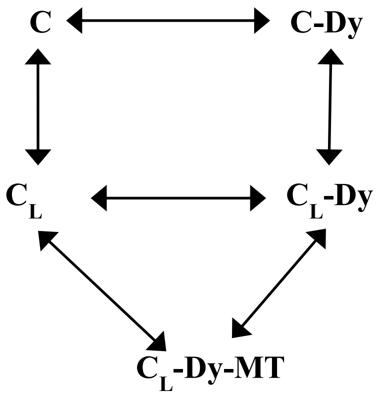


Our results suggest that the copy number of dynein involved in transport does not vary with aggregate size (Fig. 3F). For simplicity, we let the dynein copy number be one. Therefore, the cargo, once forming the active transport complex, is transported at an intrinsic velocity $v_{\mathrm{C}} = \frac{F_{\mathrm{s}}}{6\pi\eta R}$ ($F_{\mathrm{s}}$ is the dynein stalling force, $R = d\ /\ 2$ is the cargo radius and $d$ is the diameter, $\eta$ is the dynamic viscosity of the cytosol), as defined by Stokes' law[29] (Fig. 3F).

The average transport velocity $v_0$ can be written as the product of the probability of cargo forming the active transport complex $P_{\mathrm{C_L-Dy-MT}}$ and the intrinsic velocity $v_{\mathrm{c}}$:

$$v_0 = P_{\mathrm{C_L-Dy-MT}} \times v_{\mathrm{c}}.$$

We next derive how $P_{\mathrm{C_L-Dy-MT}}$ depends on system parameters. Under the equilibrium assumption, the likelihood of cargo in each state is determined by the system partition function $Z_i$ ($i$ = cargo states):

$$P_{\mathrm{C_L-Dy-MT}} = \frac{Z_{\mathrm{C_L-Dy-MT}}}{Z_{\mathrm{C_L-Dy-MT}} + Z_{\mathrm{C}} + Z_{\mathrm{C_L}} + Z_{\mathrm{C-Dy}} + Z_{\mathrm{C_L-Dy}}}.$$

To calculate individual partition functions, we model cargo as a sphere and dynein as a rod that only constrains the distance between cargo and microtubule but does not interfere with other degrees of freedom (DOFs). For each term:

**1) $Z_{\mathrm{C_L-Dy-MT}}$**

$$Z_{\mathrm{C_L-Dy-MT}} = Z_{\mathrm{C_L-Dy-MT,rot}} \times Z_{\mathrm{C_L-Dy-MT,spin}} \times e^{\frac{-E_{\mathrm{C}}-E_{\mathrm{MT}}}{kT}} \times N_r,$$

where $Z_{\mathrm{C_L-Dy-MT,rot}}$ is the rotational partition function for the cargo around the Dy-MT joint, $Z_{\mathrm{C_L-Dy-MT,spin}}$ is the spin partition function; $N_r$ is the number of dynein adaptors on cargo, proportional to the surface area $4\pi R^2$. $E_{\mathrm{C}}$ and $E_{\mathrm{MT}}$ are the binding energies between cargo and dynein or between dynein and microtubule. $Z_{\mathrm{C_L-Dy-MT}}$ can be expanded as

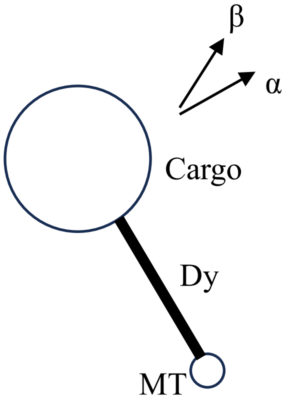


$$Z_{\mathrm{C_L-Dy-MT}} = e^{\frac{-E_{\mathrm{C}}-E_{\mathrm{MT}}}{kT}} \times N_r \times \frac{1}{h^2}\int\int e^{\frac{\left(-p_\alpha^2-p_\beta^2\right)}{2IkT}} dp_\alpha dp_\beta d\alpha d\beta \times Z_{\mathrm{spin}}$$

$$= e^{\frac{-E_{\mathrm{C}}-E_{\mathrm{MT}}}{kT}} \times 4\pi R^2\sigma \times \frac{1}{h^2} \times 4\pi^2 \int\int e^{\frac{\left(-p_\alpha^2-p_\beta^2\right)}{2IkT}} dp_\alpha dp_\beta \times Z_{\mathrm{spin}}$$

$$= e^{\frac{-E_{\mathrm{C}}-E_{\mathrm{MT}}}{kT}} \times 4\pi R^2\sigma \times \frac{1}{h^2} \times 4\pi^2 \times \sqrt{2\pi IkT}^{2} \times Z_{\mathrm{spin}}$$

$$= e^{\frac{-E_{\mathrm{C}}-E_{\mathrm{MT}}}{kT}} \times \frac{32\pi^4\sigma kT}{h^2} \times R^2 \times \frac{4}{3}\pi\rho\left(\frac{7}{5}R^5 + 2R^4 l_{\mathrm{Dy}} + R^3 l_{\mathrm{Dy}}^2\right) \times Z_{\mathrm{spin}}$$

$$= e^{\frac{-E_{\mathrm{C}}-E_{\mathrm{MT}}}{kT}} \times \frac{128\pi^5\sigma\rho kT}{3h^2} \times Z_{\mathrm{spin}} \times \left(\frac{7}{5}R^7 + 2R^6 l_{\mathrm{Dy}} + R^5 l_{\mathrm{Dy}}^2\right),$$

where $\alpha$ and $\beta$ are angular coordinates; $p_\alpha$ and $p_\beta$ are the conjugate angular momentum; $\sigma$ is the surface density of the dynein binding sites; $\rho$ is the density of the cargo; $I$ is the moment of inertia as a function of dynein length $l_{\mathrm{Dy}}$ and cargo radius $R$; $T$ is the temperature; $h$ is the Planck constant; $k$ is the Boltzmann constant.

**2) $Z_{\mathrm{C}}$**

The partition function for free cargo only contains the translational and spin DOFs.

$$Z_{\mathrm{C}} = \frac{1}{h^3}\int\int\int e^{\frac{(-p_x^2-p_y^2-p_z^2)}{2mkT}} dp_x dp_y dp_z \int\int\int dxdydz \times Z_{\mathrm{spin}}$$

$$= \frac{1}{h^3} \times V_{\mathrm{free}} \times (2\pi mkT)^{\frac{3}{2}} \times Z_{\mathrm{spin}}$$

$$= \left(\frac{8\pi^2}{3h^2}kT\right)^{3/2} \times V_{\mathrm{free}} \times Z_{\mathrm{spin}} \times R^{4.5},$$

where $V_{\mathrm{free}}$ is the integration over the space where cargo can freely diffuse.

**3) $Z_{\mathrm{C-Dy}}$**

$$Z_{\mathrm{C-Dy}} = Z_{\mathrm{C}} \times N_r = 4\pi\sigma\left(\frac{8\pi^2}{3h^2}kT\right)^{3/2} \times V_{\mathrm{free}} \times Z_{\mathrm{spin}} \times R^{6.5} \times e^{\frac{-E_{\mathrm{MT}}}{kT}},$$

given the multiplicity of dynein binding sites and the negligible size of dynein compared to the cargo.

**4) $Z_{\mathrm{C_L}}, Z_{\mathrm{C_L-Dy}}$**

Generally, it is unclear how to write down the exact partition function for the local states, since its molecular nature is not clear. As explained below, this difficulty is circumvented by introducing an empirical parameter $\alpha(R)$, the likelihood of cargo in the freely-diffusing state, which can be approximated by the measurement in Fig. 2G.

**Analytical expressions for $v_0$ as a function of $R$ can be obtained under two conditions:**

**I) The active transport complex is transient or has low stability**, where most cargoes are not forming an active transport complex with dynein-microtubule at the steady state. This condition implies $Z_{\mathrm{C}}, Z_{\mathrm{C_L}} \gg Z_{\mathrm{C-Dy}}, Z_{\mathrm{C_L-Dy}}$.

$$P_{\mathrm{C_L-Dy-MT}} \approx \frac{Z_{\mathrm{C_L-Dy-MT}}}{Z_{\mathrm{C}} + Z_{\mathrm{C_L}}} = \frac{Z_{\mathrm{C_L-Dy-MT}}}{\frac{Z_{\mathrm{C}}}{\alpha(R)}},$$

$\alpha(R)$ is the is the probability of cargo in the freely-diffusing state. Under the low-stability condition, $\alpha(R) = \frac{Z_{\mathrm{C}}}{Z_{\mathrm{C}} + Z_{\mathrm{C_L}}}$. We can rewrite the above equation as

$$P_{\mathrm{C_L-Dy-MT}} \propto \left(\frac{7}{5}R^{2.5} + 2R^{1.5}l_{\mathrm{Dy}} + R^{0.5}l_{\mathrm{Dy}}^2\right) \times \alpha(R),$$

and,

$$v_0 \propto \left(\frac{7}{5}R^{2.5} + 2R^{1.5}l_{\mathrm{Dy}} + R^{0.5}l_{\mathrm{Dy}}^2\right) \times \alpha(R) \times \frac{F_{\mathrm{s}}}{6\pi\eta R}.$$

Considering $R \gg l_{Dy}$ in most examples, we get

$$v_0 \propto R^{1.5} \times \alpha(R).$$

**II) The active transport complex has high stability**, where most cargoes in the local state are engaged in the active transport complex. This condition implies $Z_{\mathrm{C-Dy}}, Z_{\mathrm{C_L-Dy}} \ll Z_{\mathrm{C_L-Dy-MT}}$.

$$P_{\mathrm{C_L-Dy-MT}} \approx \frac{Z_{\mathrm{C_L-Dy-MT}}}{Z_{\mathrm{C_L-Dy-MT}} + Z_{\mathrm{C-Dy}} + Z_{\mathrm{C}}}.$$

Under this high-stability condition, $\alpha(R) = \frac{Z_{\mathrm{C-Dy}} + Z_{\mathrm{C}}}{Z_{\mathrm{C_L-Dy-MT}} + Z_{\mathrm{C-Dy}} + Z_{\mathrm{C}}}$, therefore,

$$P_{\mathrm{C_L-Dy-MT}} \approx 1 - \alpha(R).$$

And,

$$v_0 \propto \frac{F_{\mathrm{s}}}{6\pi\eta R} \times (1 - \alpha(R)).$$
