## Supplementary Figures for "Episodic Transport of Protein Aggregates Achieves a Positive Size Selectivity in Aggresome Formation"

### Supplementary Figures (Fig. S1-10)

#### Fig. S1

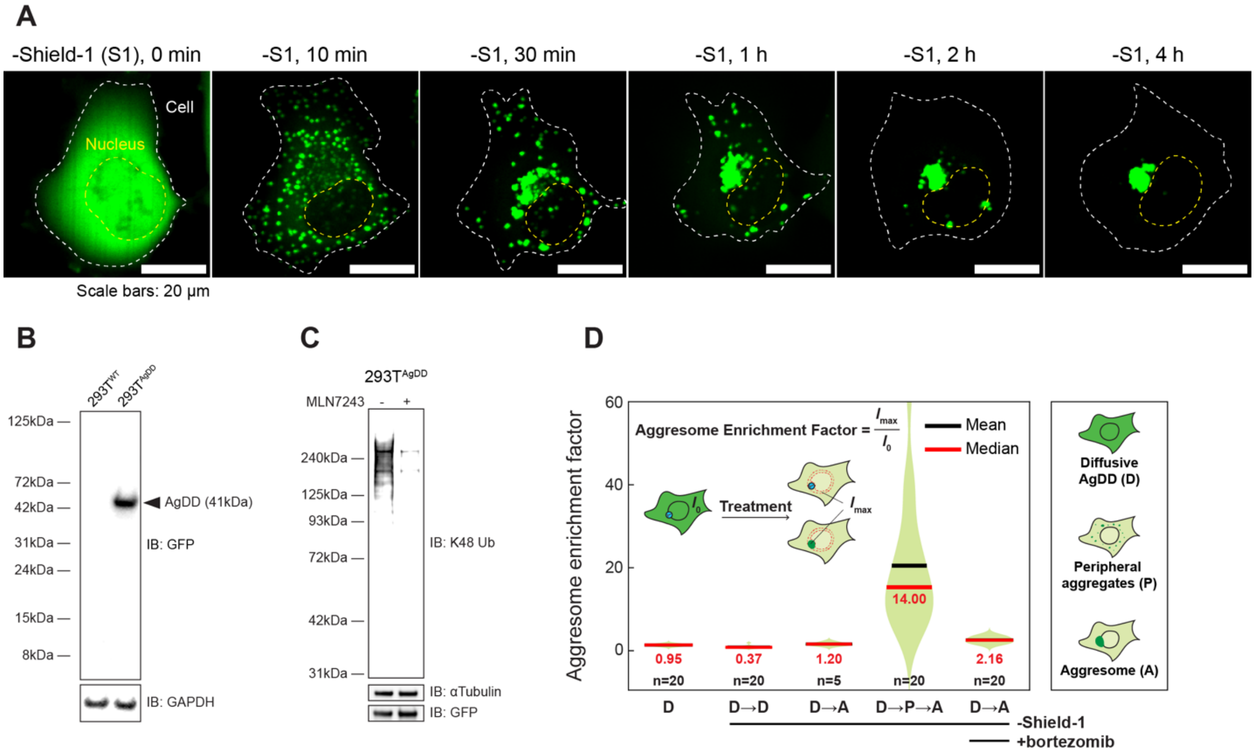


**S1. AgDD aggresome formation in cultured cells**

**(A)** Live-cell confocal images of aggresome formation in AgDD-expressing U2OS cells after Shield-1 (S1) removal. Cell and nucleus boundaries are marked by white and yellow dashed lines, respectively.

**(B)** AgDD is expressed as a full-length protein. WT or AgDD-expressing HEK293T cells were lysed and immunoblotted with an anti-GFP antibody (Proteintech, #50430-2-AP), and then re-probed with anti-GAPDH antibody (Proteintech, #10494-1-AP) as the loading control.

**(C)** Anti-K48 ubiquitin chain western blotting of AgDD-expressing HEK293T cells treated or not treated with MLN7243 (3 µM, for 3 h). S1 was removed 1 h after adding MLN7243. Cells were lysed and immunoblotted with an antibody for Lys-48-linked ubiquitin chains (Sigma, #05-1307). The blot was re-probed with anti-alpha tubulin (Novus Biologicals, #NB100-1639) and anti-GFP antibodies as controls.

**(D)** The aggresome enrichment factor was determined as the maximal fluorescent intensity of pixels within the perinuclear region ($\boldsymbol{I}_{max}$) divided by the mean pixel intensity of the whole cell ($\boldsymbol{I}_0$) before S1 removal (see Methods). Shield-1 removal and bortezomib treatment were indicated below the graph. D: diffusive AgDD signal; P: peripheral AgDD aggregates; A: AgDD aggresome. n: number of cells analyzed.

Fig. S2

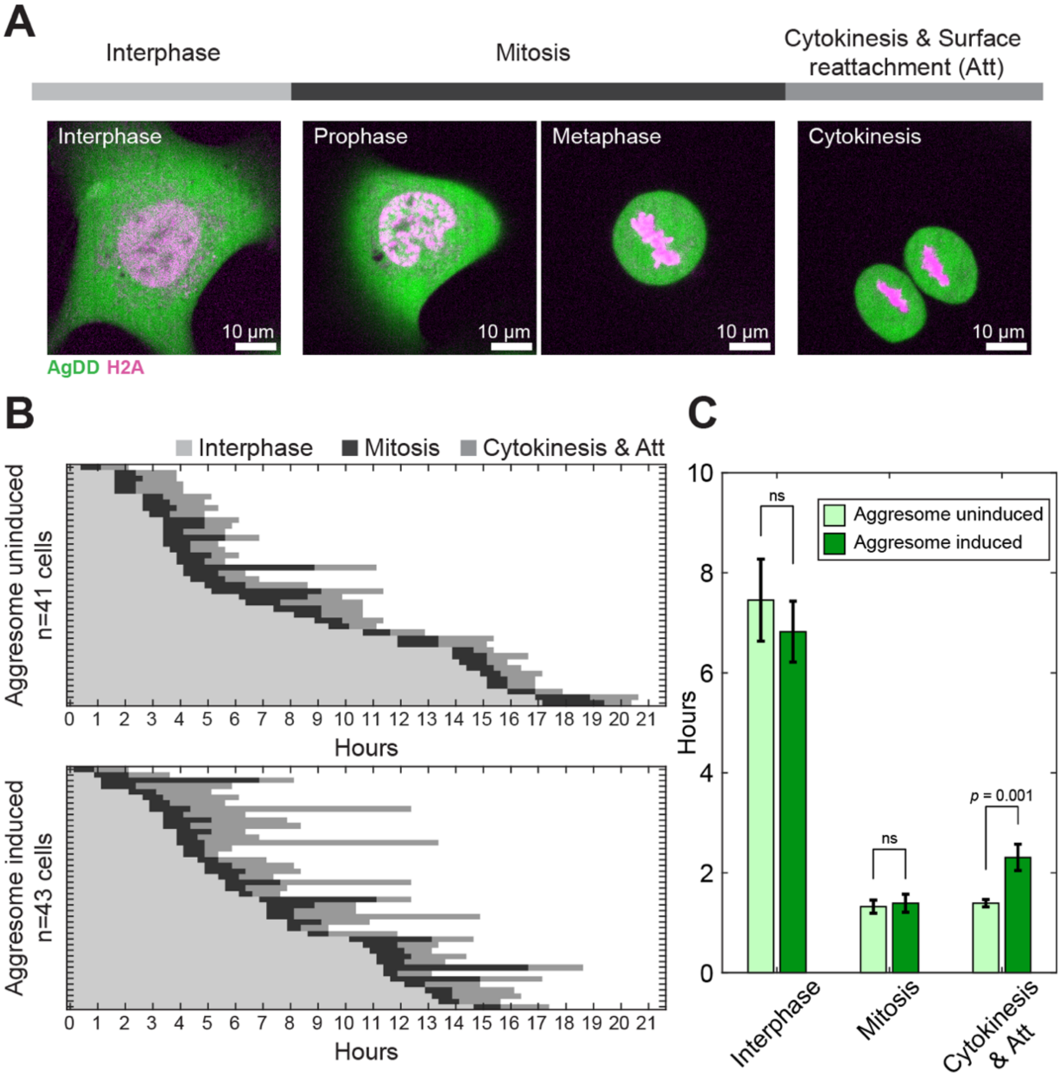


**S2. Cell cycle progression is not affected by AgDD aggresome**

**(A)** Representative live-cell confocal images showing the typical cell cycle progression of U2OS cells stably expressing AgDD (green) and HaloTag-Histone H2A (stained by JFX650-HaloTag ligand, magenta). The cell cycle was divided into three phases based on cell morphology and histone localization.

**(B)** Collection of cell cycle annotations (defined in **A**) of individual U2OS cells with or without aggresome induction. Cells were first induced by Shield-1 removal for 12 hours ("Aggresome induced") or left untreated ("Aggresome uninduced") before being imaged by a spinning-disk confocal microscope. n cells that divided once during the time-lapse were randomly selected from 5 fields of view, and the duration of each cell cycle phase is indicated by horizontal bars, colored as in **A**. Cells are ranked along the *y*-axis based on the total tracked time.

**(C)** Quantification of the duration of each cell cycle phase from **B**, presented as mean ± SEM. P-values were determined by unpaired two-tailed Student's *t*-test (ns: $p > 0.5$).

#### Fig. S3

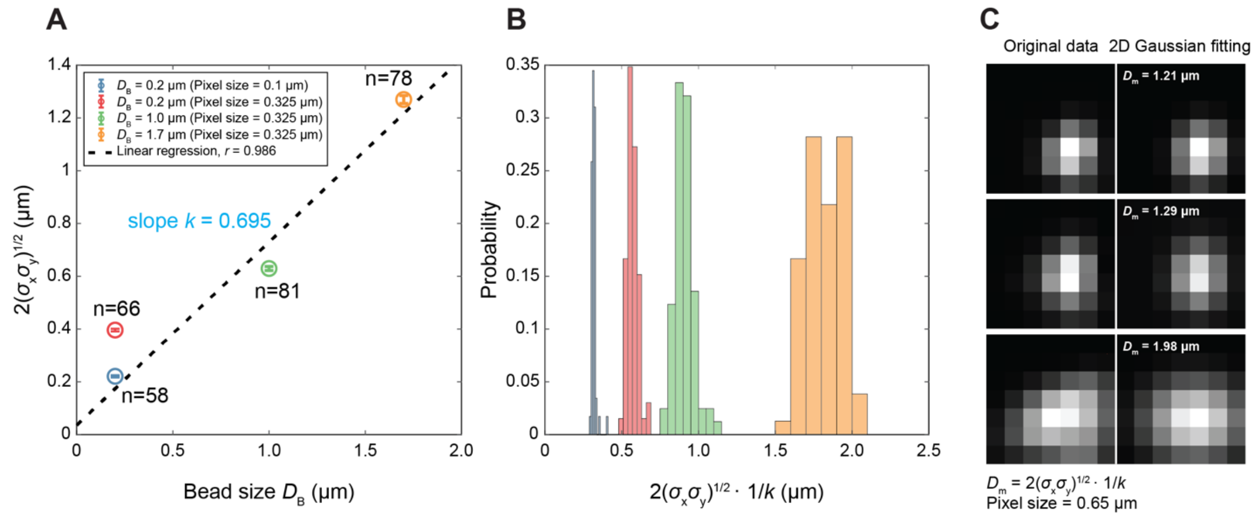


**S3. Calibration of the aggregate-size determination using fluorescent beads as size standards**

**(A)** Calibration curve using Fluoresbrite® YG Carboxylate Microspheres with different diameters ($D_B$). Beads were imaged using different magnifications indicated by the pixel size and fitted by a 2D Gaussian function with the spread parameters $\sigma_X$ and $\sigma_Y$ along the $x$ and $y$-axis (see Methods). Linear regression was performed on the nominal size of beads and $2\sqrt{\sigma_X\sigma_Y}$ to generate a cofactor $1/k$ that will be used to convert the experimentally determined spread parameters to the particle diameter, as shown in **B**. Data points in **A** show mean ± SEM. n: number of beads analyzed.

**(C)** Example images of individual AgDD aggregates (left) and the best 2D Gaussian fitting (right). $D_m$: measured diameter of the aggregate.

#### Fig. S4

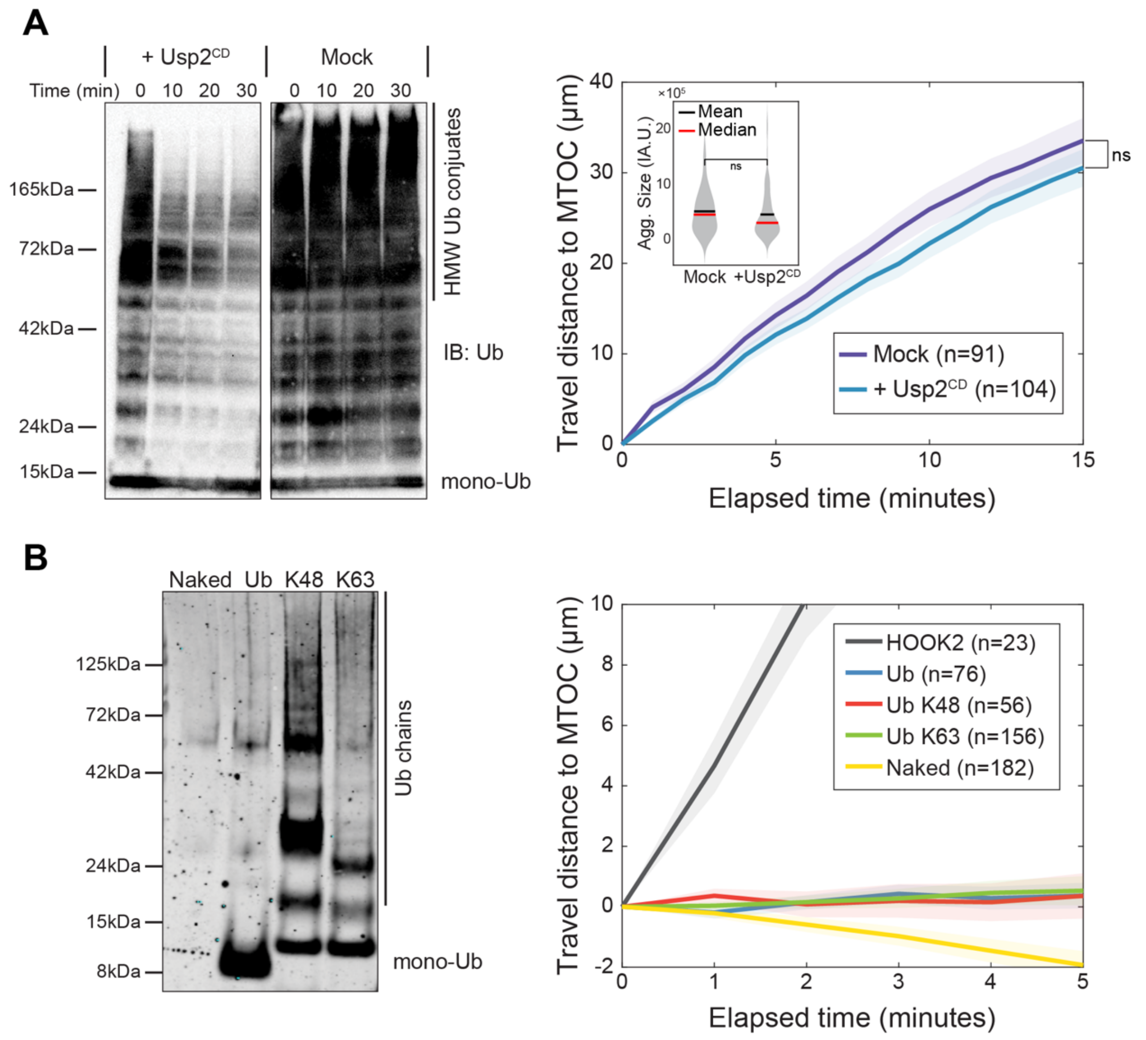


**S4. Ubiquitylation is not required for AgDD-mediated aggresome formation in XE(A)** Usp2 catalytic domain (Usp2$^{CD}$) did not perturb aggregate transport in XE. Left: Interphase XE was incubated with or without 10 µM Usp2$^{CD}$ at 18 °C for different times and immunoblotted using an anti-ubiquitin (Ub) antibody (Cell Signaling, #43124S). HMW: high molecular weight. Right: AgDD aggregates pretreated with or without 10 µM Usp2$^{CD}$ for 10 minutes were tested for mobility in XE. Data shown are the average trajectories of aggregates plotted over time with mean (thick line) ± SEM (shading). n: number of trajectories. P-value was determined for the final positions of aggregates under two conditions at $t$ = 15 min using unpaired two-tailed Student's $t$-test (ns: $p = 0.35$). Inset: distributions of aggregates' size indicated by the total fluorescence intensity of each particle; P-value was determined by unpaired two-tailed Student's $t$-test (ns: $p = 0.4$).

**(B)** Purified Ub chains did not support bead transport in XE. Left: Ub or Ub chains of indicated linkages (top) were synthesized, biotinylated and conjugated to Dynabeads through biotin-tags as described in Methods. Dynabeads were passivated with PEG polymers before conjugation to reduce nonspecific binding or transport (Methods). Conjugated-bead samples were validated by heating the beads with SDS sample buffer and blotting with Alexa Fluor™ 790 streptavidin (ThermoFisher, #S11378). Right: average trajectories of beads conjugated with indicated Ub construct. Unconjugated but passivated beads ("Naked") and HOOK2-coated beads were included for comparison. n: number of beads analyzed.

**Fig. S5**

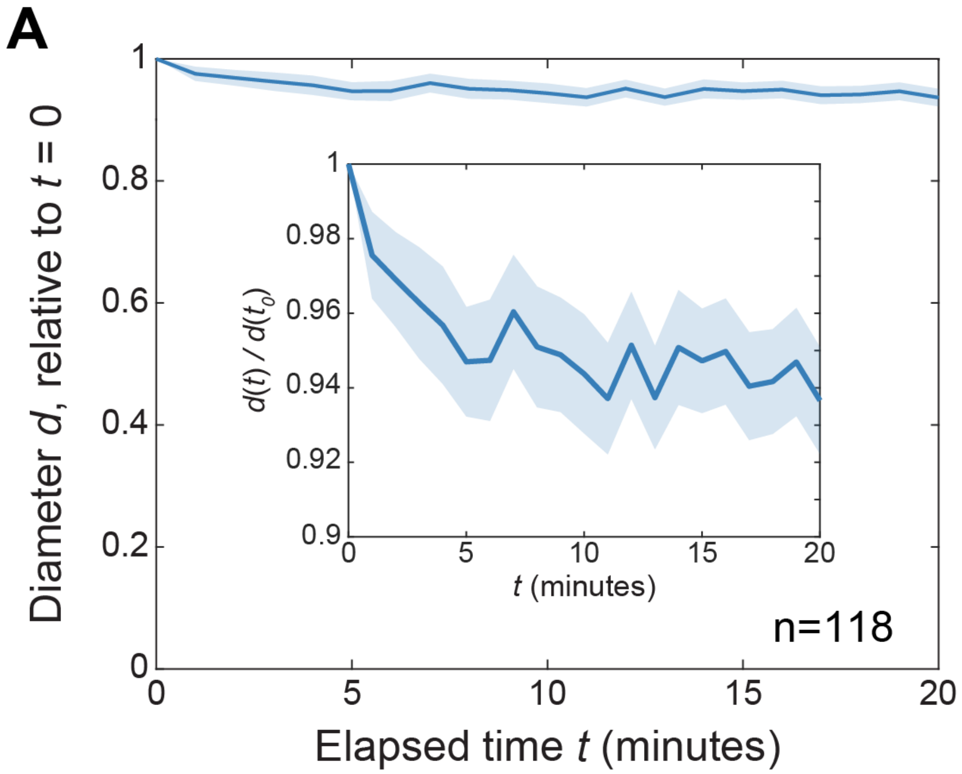


**S5. AgDD aggregates' diameter does not change significantly during the transport assay in XE**

The mean of the aggregate's diameter relative to the initial value at $t = 0$ is plotted as mean (thick line) ± SEM (shading). n = 118 aggregates from the experiment in **Fig. 2D** that can be tracked for at least 20 minutes were used for analysis. Inset: a view with local *y*-axis.

### Fig. S6

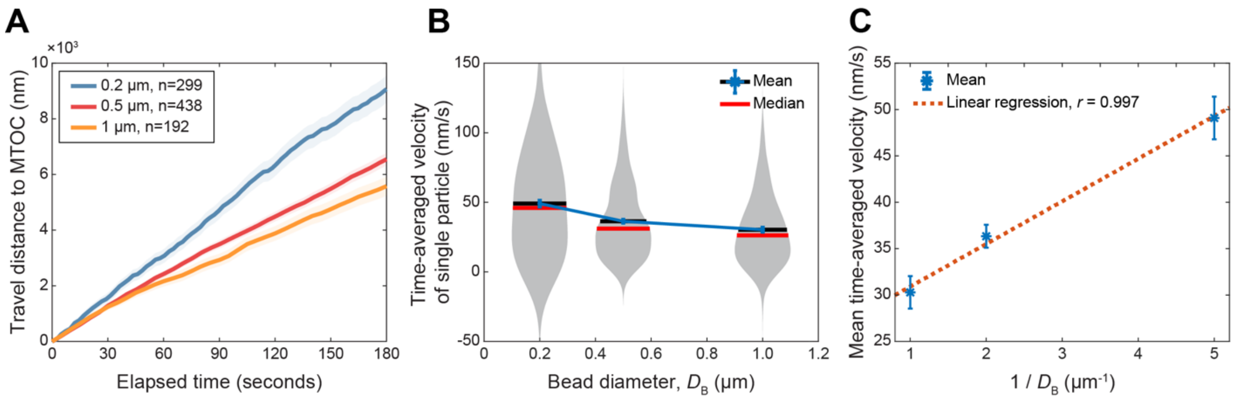


**S6. Transport of Fluoresbrite® YG Carboxylate Microspheres in XE shows a negative size selectivity**

**(A)** Averaged trajectories of unpassivated beads with different diameters in XE, plotted as mean (thick line) ± SEM (shading). Data were acquired as described in **Fig. 2A**, but at 12 frames per minute for 20 minutes. The time-averaged velocities of individual particles were calculated as in **Fig. 2D** and presented in **B**. The mean values are plotted against the inverse of bead's diameter in **C**. n: number of trajectories.

**Fig. S7**

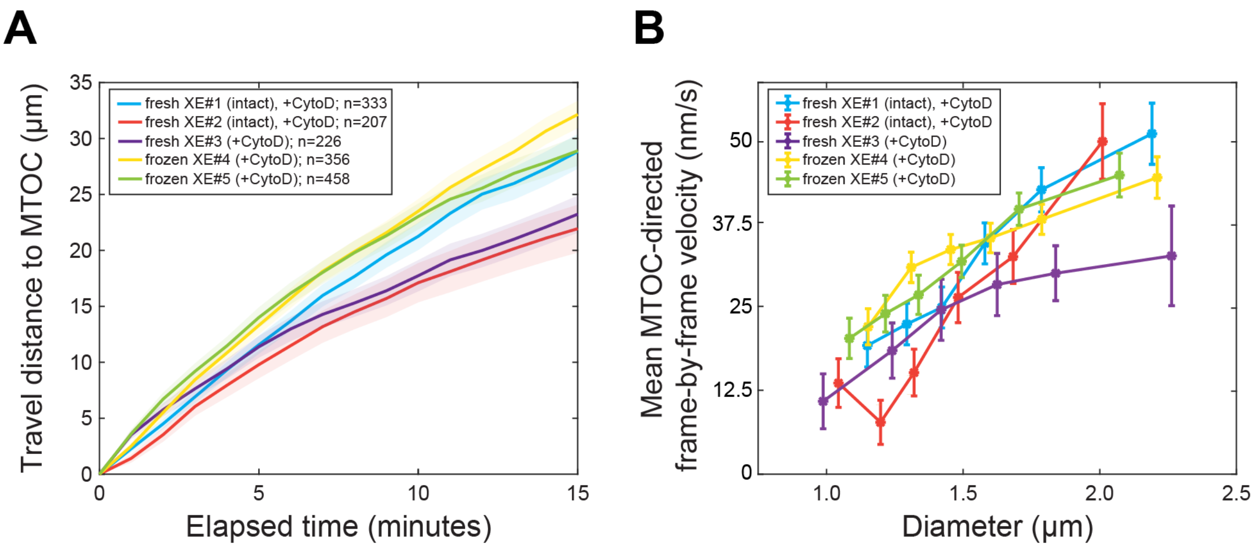


**S7. Reproducibility of aggregate transport assay in different batches of XE**
**(A)** Averaged trajectories of AgDD aggregates in XE prepared freshly (XE#1-3) or from frozen stock (XE#4,5); Cytochalasin D (CytoD) was added to the XE during preparation (XE#3-5) or right before the experiment (XE#1,2; see Methods). Results are plotted as the mean (thick line) ± SEM (shading). n: number of trajectories. The same data were used to determine the size dependence of the MTOC-directed frame-by-frame velocity as described in **Fig. 1J** and presented in **B** as mean ± SEM. XE#1 and XE#3 were used in the experiments in **Fig. 2**.

**Fig. S8**

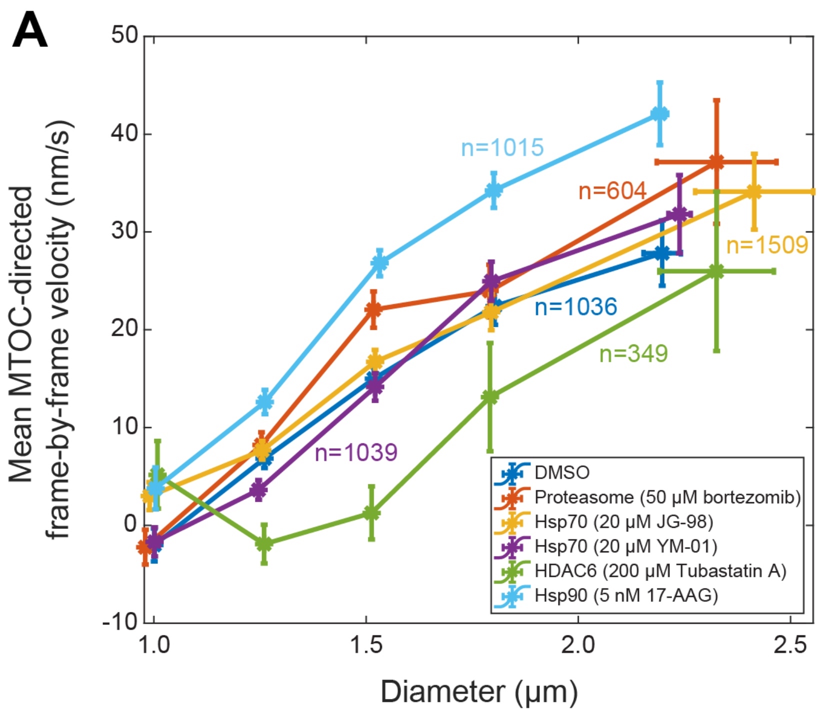


**S8. HDAC6 inhibitor reduces aggregate transport velocity and increases the size threshold for transport**

MTOC-directed frame-by-frame velocities of AgDD aggregates in XE treated with indicated inhibitors were determined as described in **Fig. 1J** and grouped by the aggregates' diameter. The plot shows the mean value within each size group ± SEM. Data were acquired once per minute for 30 minutes. n: number of trajectories.

#### Fig. S9

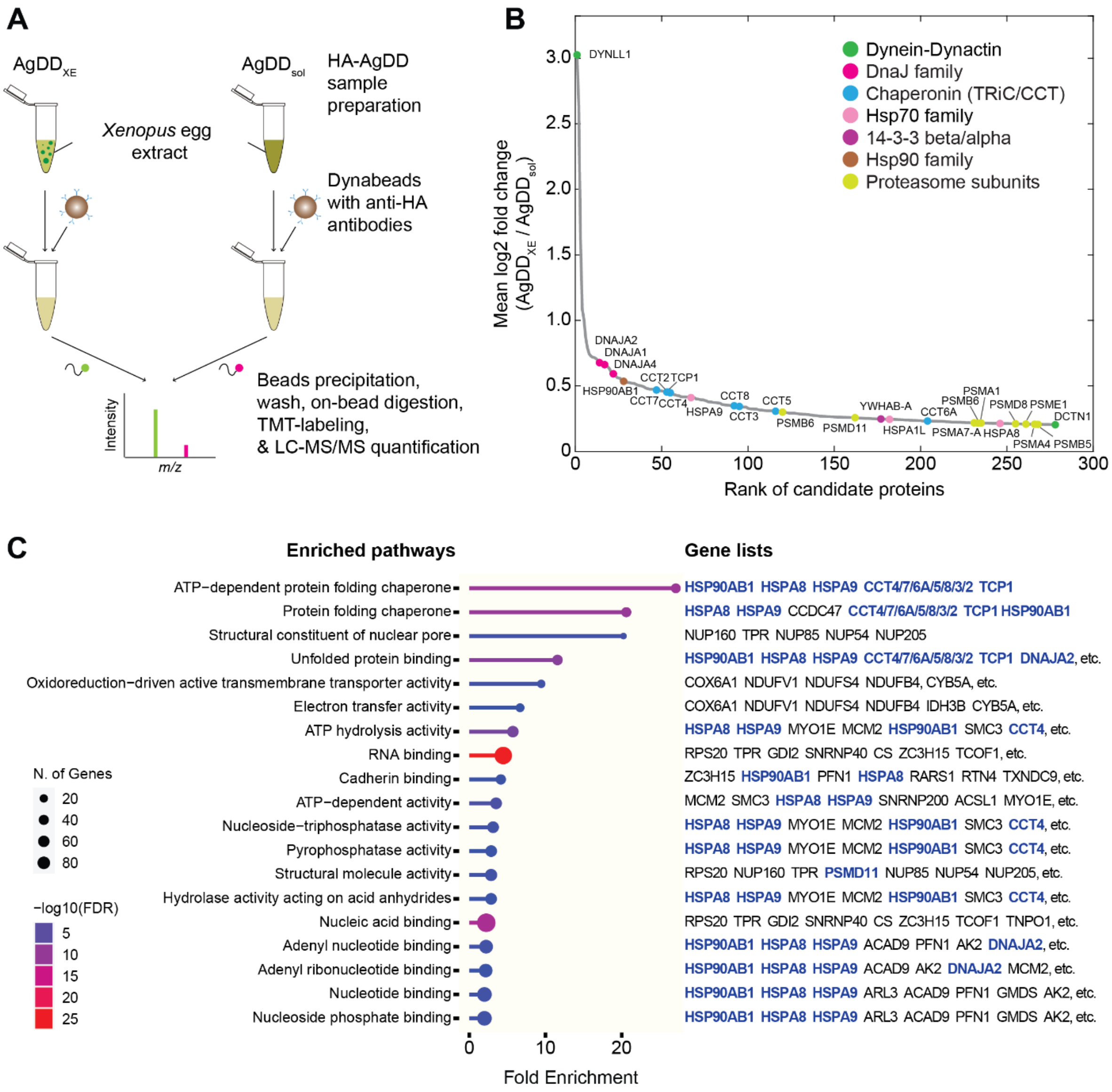


**S9. Identification of the interactome of AgDD aggregates in XE**

**(A)** Schematic of the workflow of immunoprecipitation-based mass spectrometry (IP-MS) using Dynabeads coated with AgDD aggregates formed in XE ($AgDD_{XE}$) and soluble AgDD stabilized by Shield-1 ($AgDD_{sol}$). AgDD contains an HA-tag at the N terminus for bead conjugation. The relative abundance of proteins captured by different beads was quantified by Tandem-Mass-Tag mass spectrometry (TMT-MS) (see Methods).

**(B)** Identified proteins are ranked by the fold change ($AgDD_{XE}/AgDD_{sol}$). Proteins of interest are color-coded and labeled as indicated in the legend.

**(C)** Gene Ontology (GO) analysis of the top 280 hits in **B** by shinyGo (see Methods). Part of the gene list associated with each GO term is shown on the right, and the genes that belong to the protein families listed in **B** are highlighted in blue. The complete gene lists are in the **Supplementary Table S1**.

Fig. S10

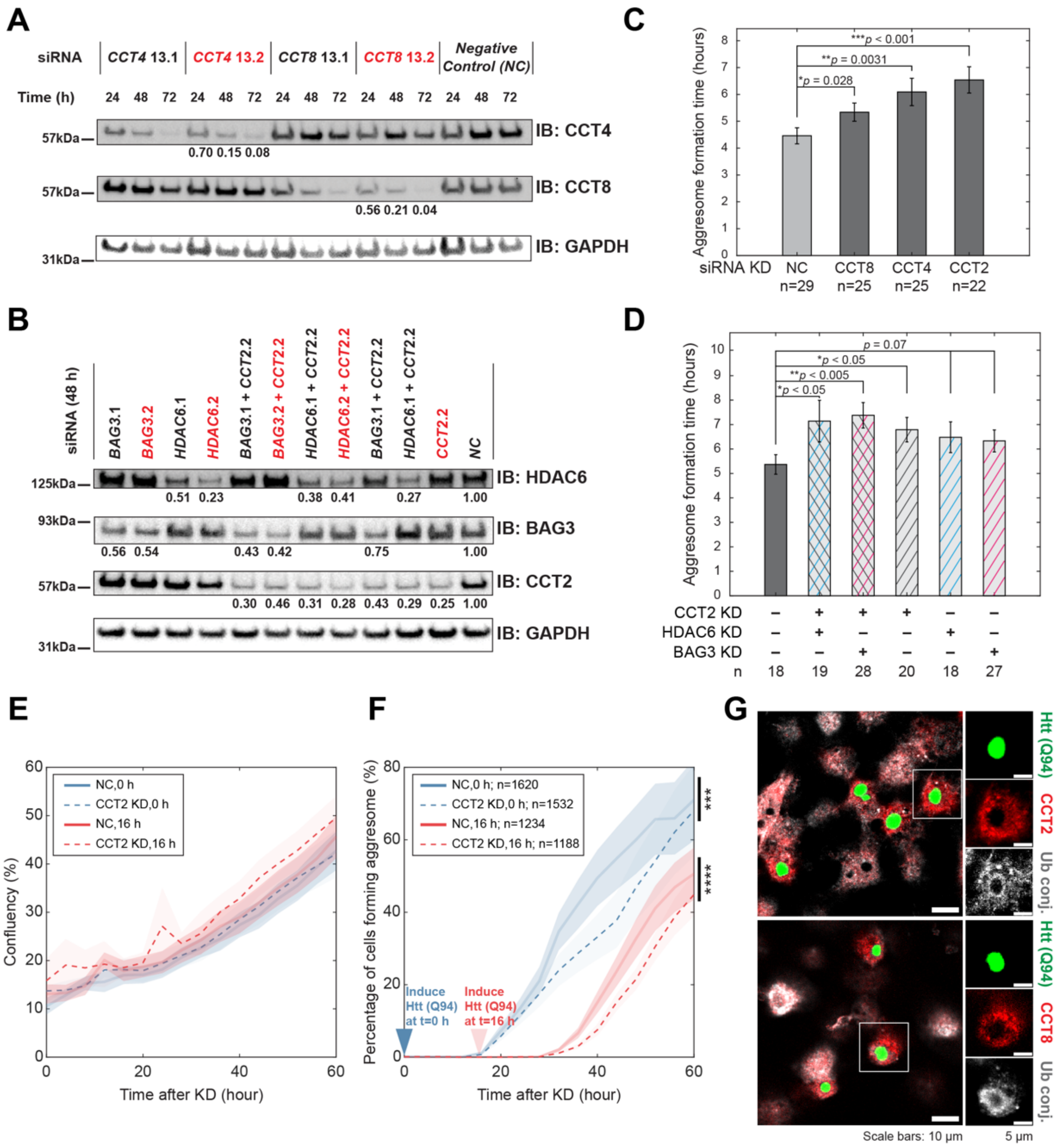


**S10. Chaperonin components play a role in aggresome formation**
**(A)(B)** Single and double knock-down (KD) of chaperonin subunits CCT2, CCT4, CCT8, and BAG3, HDAC6 in HEK293T cells. KD efficiency was examined by immunoblotting against each target protein and labeled below the band as the ratio of the protein in KD cells versus in the negative control (NC, transfected with the scrambled RNA). siRNAs used for the experiments in **C** and **D** are highlighted in red.

**(C)(D)** Effects of gene KD on AgDD aggresome formation. AgDD aggresome formation was induced in cells 48 hours after siRNA transfection via Shield-1 removal. Cells were analyzed by live-cell microscopy and the time from the appearance of peripheral aggregates to the complete sequestration of all aggregates into the single perinuclear punctum of aggresome was determined on randomly selected (n) cells. Bar plot shows the mean ± SEM. P-values were calculated using an unpaired one-tailed Student's *t*-test.

**(E)(F)** Chaperonin KD delayed aggresome formation by poly-glutamine protein Htt (Q94). Expression of Htt (Q94)-CFP from a TetOn promoter in HEK293T cells was induced by adding 1 μg/mL doxycycline together with or 16 hours after transfection of the CCT2 siRNA (t=0, 16 h). Cells were imaged by IncuCyte® ZOOM Live-Cell Analysis System for 60 hours. No significant growth defects were detected after CCT2 KD or Htt (Q94) expression during this period, as indicated by the growth curves in **E**. The formation of poly-Q aggresome was determined at each time point as described in Methods and presented in **F**. n: number of cells with an aggresome at the end of the experiment. Curves show mean ± SEM (over 9 fields of view). P-values were determined by one-sample *t*-test at the 5% significance level, given the null hypothesis that the mean of $(f_{\mathrm{KD}} - f_{\mathrm{NC}})/f_{\mathrm{NC}}$ of all time points after adding doxycycline is zero, where $f$ is the percentage of cells with an aggresome (*y*-axis of **F**) (***$p$ = 0.0004; ****$p$ = 0.0001).

(G) Example immunofluorescence images of Htt (Q94)-expressing (green) HEK293T cells stained with anti-CCT2 (red, above), anti-CCT8 (red, bottom), and anti-ubiquitin conjugates (Ub conj., gray) antibodies. WT cells were transfected with pTetOn-Htt (Q94)-CFP, incubated for 14 hours and then induced with 1 μg/mL doxycycline for 12 hours before fixation.

### Supplementary Videos (Supp. Video 1-8)

**SV1. Aggresome formation in U2OS cells stably expressing AgDD**
Aggresome formation in U2OS cells stably expressing AgDD after Shield-1 removal, imaged at 60x every 1 minute by a spinning-disk confocal microscope.

**SV2. Aggregation of purified AgDD in interphase XE**
Aggregation of purified AgDD was induced in interphase *Xenopus* egg extract (XE) at 18 °C by adding a 20-fold molar excess of recombinant FKBP(F36V) to rapidly deplete free Shield-1, imaged at 10x every 1 minute by a widefield fluorescent microscope.

**SV3. Transport of AgDD aggregates in interphase XE**
Transport of AgDD aggregates in interphase XE, imaged at 10x every 1 minute by a widefield fluorescent microscope. The microtubule aster was labeled with tubulin-Alexa647. Actin was depolymerized by cytochalasin D.

**SV4. Transport of AgDD aggregates in cycling XE with intact actin**
Transport of AgDD aggregates in cycling XE with intact actin, imaged at 10x every 1 minute by a spinning-disk confocal microscope. The MTOC was labeled with EB1-mApple. Image acquisition was started 15 minutes after warming up the XE to 18 °C, when the XE was in interphase.

**SV5. CC1 inhibits the transport of AgDD aggregates in interphase XE**
CC1 inhibited the transport of AgDD aggregates in interphase XE. The experiment was performed as in SV3, but with 40 μg/mL CC1 in the XE.

**SV6. AgDD aggregates formed in a buffer could not be transported towards the MTOC**
AgDD aggregates formed in a buffer (1 mM dithiothreitol, 100 mM $KCH_3COOH$, 30 mM KCl, 1 mM $MgCl_2$, 1 mM $Na_2ATP$, 10 mM $Na_2HPO_4$) could not be transported towards the MTOC. The experiment was performed as in **SV3**.

**SV7. Transport of Dynabeads coated with chaperonin in interphase XE**
Transport of Dynabeads coated with chaperonin in interphase XE, same data as shown in **Fig. 4A**. The experiment was performed as in **SV3**.

**SV8. Aggresome formation in U2OS cells stably expressing AgDD-HOOK2**
Aggresome formation in U2OS cells stably expressing AgDD-HOOK2 after Shield-1 removal, imaged at 60x every 1 minute by a spinning-disk confocal microscope. Three cells were shown in the movie: the top right one spontaneously forming aggresome before Shield-1 removal, the top left one forming aggresome without peripheral aggregates, and the middle one forming distinguishable peripheral aggregates before aggresome formation.

#### Supplementary Tables

**Table S1. Complete gene lists of GO terms enriched by shinyGo**
Complete gene lists of GO terms enriched by shinyGo, derived from the interactome of AgDD aggregates (**Fig. S9**; Methods).

**Table S2. TMT-MS quantification of the designated antigen amount on the beads**
TMT-MS quantification of the designated antigen amount on the beads coated with antibodies for different factors and dynein adaptors (**Fig. 4**; Methods).

#### Supplementary Data Files

**Supp. Data File 1. TMT-MS quantification of AgDD interactomes**
TMT-MS quantification of AgDD interactomes. Both raw data and the identified interactome of AgDD aggregates used for GO term analysis are included.

**Supp. Data File 2. TMT-MS quantification of adaptor-coated beads**
Raw data from TMT-MS quantification of beads coated with different factors are included.
